# Gene-environment interactions characterized by single embryo transcriptomics

**DOI:** 10.1101/805556

**Authors:** Alfire Sidik, Groves B. Dixon, Hannah G. Kirby, Johann K. Eberhart

**Affiliations:** Department of Molecular Biosciences, Institute for Cellular and Molecular Biology, Institute for Neuroscience, Waggoner Center for Alcohol and Addiction Research, University of Texas, Austin, TX 78712, USA

**Author notes:** Correspondence to: Johann K. Eberhart, PAT 522, The University of Texas at Austin, Molecular Biosciences, College of Natural Sciences, 2401 Speedway, Austin, TX, 78712.

## Abstract

Gene-environment interactions are likely to underlie most human birth defects. The most common environmental contributor to birth defects is likely prenatal alcohol exposure. Fetal Alcohol Spectrum Disorders (FASD) describes the full range of defects that result from prenatal alcohol exposure. Gene-ethanol interactions underlie susceptibility to FASD but we lack a mechanistic understanding of these interactions. Here, we leverage the genetic tractability of zebrafish to address this problem. We first show that *vangl2*, a member of the Wnt/planar cell polarity (Wnt/PCP) pathway that mediates convergent extension movements, strongly interacts with ethanol during late blastula and early gastrula stages. Embryos mutant or heterozygous for *vangl2* are sensitized to ethanol-induced midfacial hypoplasia. We performed single-embryo RNA-Seq during early embryonic stages, to assess individual variation to the transcriptional response to ethanol and determine the mechanism of the *vangl2*-ethanol interaction. To identify the pathway(s) that are disrupted by ethanol we used these global changes in gene expression to identify small molecules that mimic the effects of ethanol via the Library of Integrated Network-based Cellular Signatures (LINCS L1000) dataset. Surprisingly, this dataset predicted that the Sonic Hedgehog (Shh) pathway inhibitor, cyclopamine, would mimic the effects of ethanol, despite the fact that ethanol did not alter the expression levels of direct targets of Shh signaling. Indeed, we found that ethanol and cyclopamine strongly interact to disrupt midfacial development. Collectively, these results suggest that the midfacial defects in ethanol-exposed *vangl2* mutants are due to an indirect interaction between ethanol and the Shh pathway. Vangl2 functions as part of a signaling pathway that regulates coordinated cell movements during midfacial development. Consistent with an indirect model, a critical source of Shh signaling that separates the developing eye field into bilateral eyes, allowing the expansion of the midface, becomes mispositioned in ethanol-exposed *vangl2* mutants. We demonstrate that ethanol also interacts with another Wnt/PCP pathway member, *gpc4*, and a chemical inhibitor, blebbistatin. By characterizing membrane protrusions, we demonstrate that ethanol synergistically interacts with the loss of *vangl2* to disrupt cell polarity required for convergent extension movements. Collectively, our results shed light on the mechanism by which the most common teratogen can disrupt development.

## Introduction

Birth defects manifest as structural or functional malformations at birth and are the leading cause of infant mortality in the United States [1]. Although the multifactorial etiologies of birth defects are not well understood, many are thought to derive from a complex interplay between genetic and environmental factors. The adverse effects of teratogens, or environmental agents that cause irreversible developmental defects, were first recognized in the 1950s and 1960s [2]. It was then that investigators began to observe developmental anomalies in infants exposed to methylmercury and thalidomide in utero [3, 4]. Since then, other environmental teratogens have come to light, including but not limited to, pollutants, pharmaceuticals, and chemicals.

Prenatal alcohol exposure (PAE) is the most common cause of birth defects, with the prevalence of fetal alcohol spectrum disorders (FASD) in the US is as high as 2-5% [5]. Fetal alcohol syndrome (FAS) is the most severe outcome following PAE and is characterized by midfacial hypoplasia, as well as growth and neural deficits [6]. While PAE is required for the development of FASD, the teratogenic effects of ethanol are modulated by genetics [7, 8]. For instance, monozygotic twins were 100% concordant for FAS, whereas dizygotic twins were only 63% concordant [8, 9]. Furthermore, animal models of FAS show strain-specific differences after controlling for environmental variables such as dose and timing [10, 11]. Despite this, the genetic factors that protect or predispose an individual to FASD are poorly understood. Moreover, we still lack a basic understanding of the mechanism of ethanol teratogenesis. Zebrafish are well suited to address this problem due to their genetic tractability and ease of embryological manipulation [12].

In a screen to identify genetic modifiers of ethanol teratogenicity, *vang-like 2* (*vangl2*), a member of the non-canonical Wnt/PCP pathway, emerged as an ethanol sensitive locus [13]. In zebrafish, mutations in the *vangl2* locus disrupt convergent extension movements during gastrulation, resulting in a shortened, broadened, body axis [14]. These mutants infrequently present with cyclopia, fusion of the bilateral eyes, but the phenotypic expressivity of this trait varies with factors such as temperature and genetic background [15]. In the screen, all untreated *vangl2* mutants displayed proper separation of the eyes and craniofacial skeletal elements were intact [13]. Upon exposure to a subteratogenic dose of ethanol, *vangl2* mutants were fully penetrant for cyclopia and displayed profound defects to the midfacial skeleton [13]. Ethanol-treated *vangl2* heterozygotes were largely indistinguishable from their wild-type siblings, with the exception of a single synophthalmic ethanol-treated heterozygote, providing evidence for latent haploinsufficiency. Together, these data suggest a synergistic interaction between a mutation in *vangl2* and ethanol.

We know *vangl2* plays a critical role in mediating convergent extension movements as evidenced by their body axis defect, however because the early effects of ethanol exposure remain poorly defined, the precise mechanism of the *vangl2*-ethanol interaction remain elusive. To better understand how ethanol interacts with loss of *vangl2* to alter phenotypic outcomes, we took an unbiased approach to assess the transcriptional response to ethanol. we performed single embryo RNA-sequencing (RNA-seq) on control (untreated) and ethanol-treated wild-type embryos in a time-course spanning gastrulation and early segmentation stages in zebrafish.

Bioinformatic and functional analyses indicated that midfacial defects in ethanol-exposed *vangl2* mutants was due to an indirect interaction between ethanol and the Shh pathway. While there was no alteration in the level of expression of direct Shh targets, a critical source of Shh signaling that separates the developing eye field into bilateral eyes becomes mispositioned in ethanol-exposed *vangl2* mutants. We demonstrate that ethanol and loss of *vangl2*, synergistically interact to disrupt polarized cellular protrusions required for proper convergent extension movements.

## Results

### Early embryogenesis is the sensitive time window for *vangl2* mutants

To determine the critical time window of ethanol sensitivity for the *vangl2* mutants, we first initiated ethanol treatment at various stages comprising late blastula to early gastrula for 24 hours. The inner lens-to-lens width was used as a measure of synophthalmia and we calculated the occurrence of cyclopia to characterize ethanol-induced teratogenesis. In control conditions, the spacing of the eyes in *vangl2* mutants mirrored that of wild-type zebrafish. Ethanol-exposed *vangl2* mutants exhibited midline defects ranging in severity from synophthalmia to cyclopia across all time points examined, but these malformations were fully penetrant for cyclopia (100% fused; n=5/5) when ethanol was applied at shield stage (6 hours post fertilization, hpf) at the onset of gastrulation (Fig 1A-B). Interestingly, heterozygotes only displayed cyclopia when ethanol was applied at high stage (3.3 hpf) (22% fused; n=4/18), a time when treating wild-type embryos with higher concentrations of ethanol causes similar defects (Fig 1B) [16]. Thus, heterozygotes and homozygotes are divergent in their time window of greatest ethanol sensitivity. This may be due to a compensatory genetic mechanism in *vangl2* heterozygotes because zygotic gene expression initiates after high stage (4 hpf) in zebrafish.

**Fig 1.**
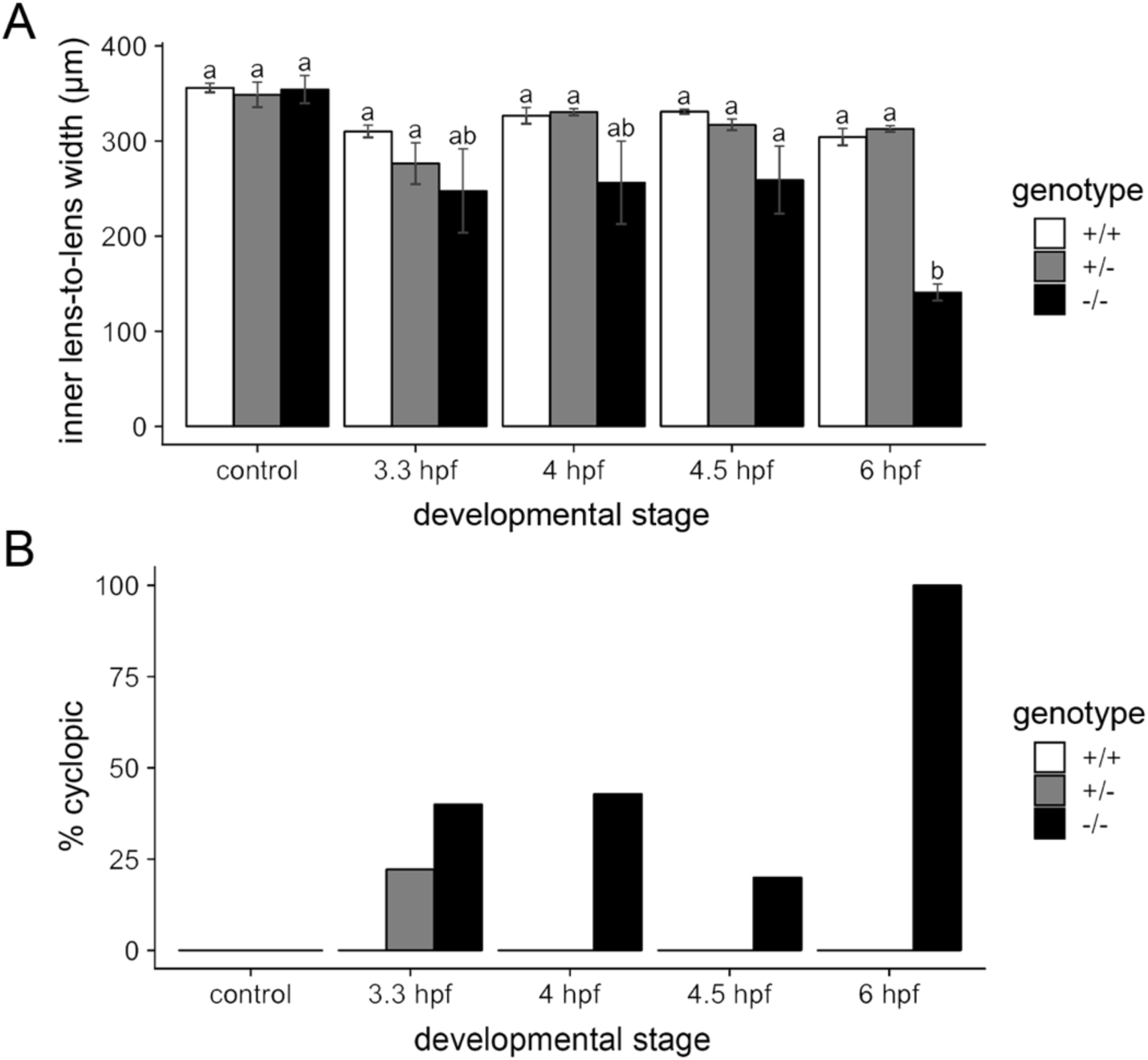
Ethanol interacts with *vangl2* during early embryogenesis. (A) Treatment of *vangl2* embryos with 1% ethanol at four different stages comprising late blastula to early gastrula [3.3, 4, 4.5, and 6 hours post fertilization (hpf)] for 24 hours. Homozygous mutants showed a significant decrease in the inner lens-to-lens width when treated from 6 to 30 hpf. (B) Heterozygous mutants were most sensitive to ethanol when exposed from 3.3 to 27.3 hpf (n=4/18 cyclopic). Homozygous mutants were most sensitive when exposed from 6 to 30 hpf (n=5/5 cyclopic).

### The effect of ethanol on the early zebrafish transcriptome is subtle relative to developmental time

To determine if ethanol caused transcriptional changes that could underlie the interaction with *vangl2*, we designed two RNA-Seq experiments that largely overlapped in design (Fig 2A-B). Embryos were exposed to a subteratogenic dose of 1% ethanol in embryo media (171 mM), which equilibrates to approximately 50 mM tissue concentration [17]. For the first experiment, embryos were treated at the onset of gastrulation (6 hpf) and collected at mid-gastrulation (8 hpf) and the end of gastrulation (10 hpf) (Fig 2A). A second experiment was performed to increase power (Fig 2B). Embryos were similarly exposed, but a 14 hpf time window was included, when the eye fields have completely separated. Each sample consisted of an individual zebrafish embryo with five replicates per timepoint and treatment. The 6 hpf control samples were omitted since they lacked ethanol-treated samples for comparison. The data from both experiments were combined for subsequent analyses, controlling for batch effects.

**Fig 2.**
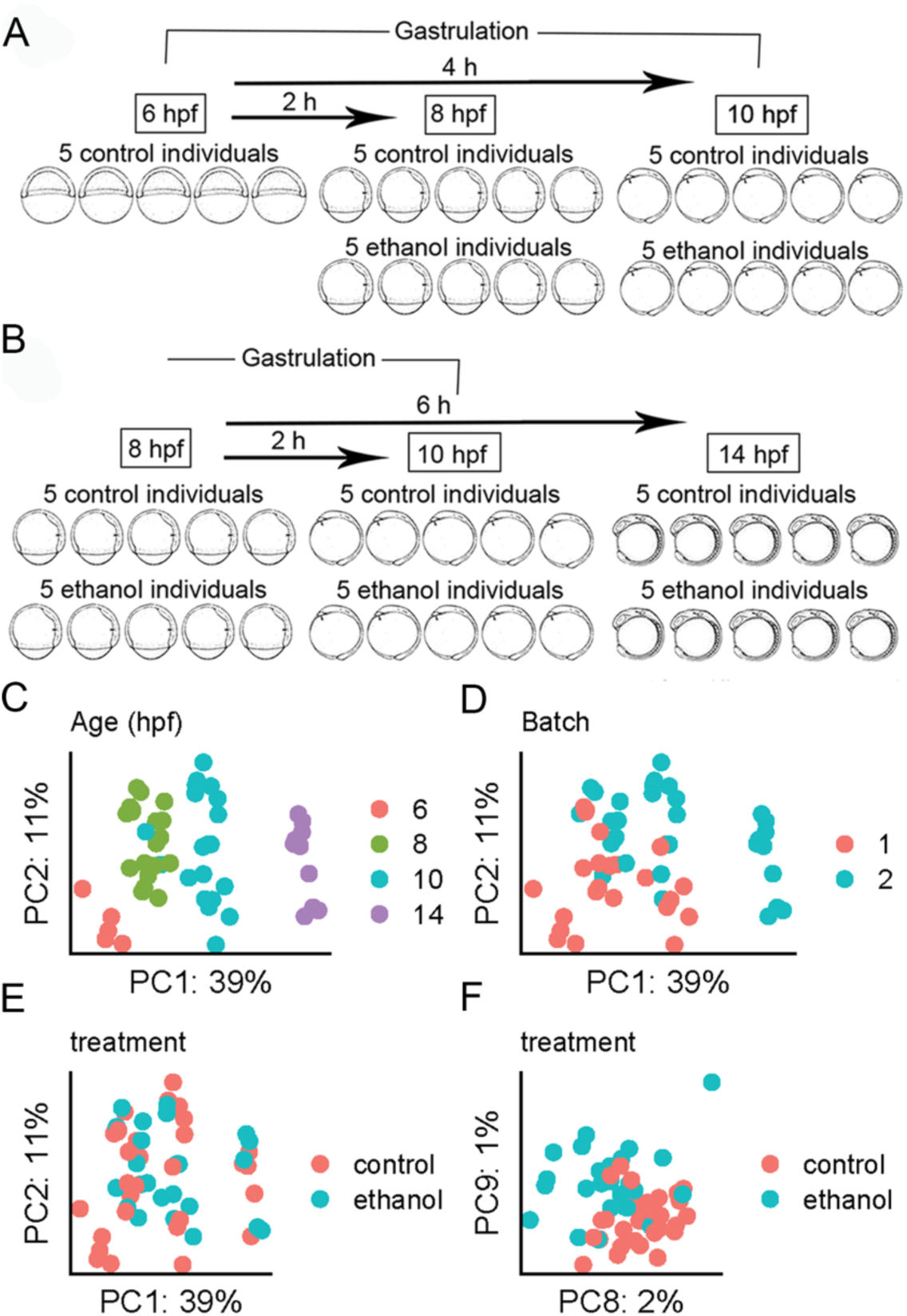
Developmental age is the dominant driver of variation in the dataset. (A) Schematic representation of the RNA-seq experimental design. Wild-type AB embryos were exposed to a subteratogenic dose of 1% ethanol in the embryo media at shield stage (6 hpf). Embryos were subsequently collected at 8 and 10 hpf for experiment 1 and (B) 8, 10, and 14 hpf, for experiment 2. Each sample consisted of a single zebrafish embryo with 5 control and 5 ethanol-treated samples per timepoint with the exception of the 6 hpf timepoint, which only had 5 control samples. (C-F) Principal components analysis (PCA) of top 25,000 most variable genes. The percentage of variance explained is given on each axis label. (C) *PC1* and *PC2* color coded by developmental timepoint. (D) *PC1* and *PC2* color coded by RNA-seq experiment (batch). (E) *PC1* and *PC2* color coded by ethanol treatment group. (F) *PC8* and *PC9* showing separation of ethanol treated and control samples.

To assess the effect of time, batch (i.e. experiment 1 and 2), and ethanol, on the early zebrafish transcriptome, we performed principal component analysis (PCA). Individuals from each timepoint clustered tightly along *PC1*, which accounted for 38% of the transcriptional variation observed across the datasets. This strong effect of time on the zebrafish transcriptome is in agreement with previous studies [18]. Clustering of samples by time, regardless of ethanol treatment suggested that control and ethanol-treated samples were accurately staged, indicating that ethanol did not delay developmentally regulated transcriptome patterns. There was greater discrimination of 14 hpf samples relative to earlier timepoints (6, 8, and 10 hpf), which is consistent with greater distinction of this timepoint in terms of developmental time and morphology (Fig 2C). *PC2* largely captured batch effects between experiment 1 and 2 (Fig 2D).

The majority of variation between samples did not appear to be due to treatment, with control and ethanol-treated samples randomly interspersed along *PC1* and *PC2* (Fig 2E). Separation by treatment was observed along *PC8* and *PC9*, which accounts for 3% of the variation in the data (Fig 2F). Hierarchical clustering of samples based on correlation, further corroborated this finding. The 6 and 14 hpf samples showed the greatest dissimilarity, whereas the 8 and 10 hpf samples showed the greatest similarity, irrespective of treatment (S1 Fig). In summary, the transcriptional effect of exposure to a subteratogenic dose of ethanol on the early zebrafish transcriptome was subtle, while time provided the strongest transcriptional fingerprint.

### Ethanol has effects on transcription that are largely distinct between different developmental timepoints

Although developmental age was a stronger source of transcriptional variation, we still detected substantial variation in gene expression following subteratogenic ethanol exposure. There were 1,414 differentially expressed genes (DEGs), with a false-discovery rate (FDR) less than 0.1 (Fig 3A; Benjamini–Hochberg procedure). There were more upregulated than downregulated DEGs among ethanol-treated individuals across timepoints (S2 Fig) and these DEGs were distinct between developmental timepoints (Fig 3A,B). In summary, while some genes are generally affected by ethanol across several critical periods, ethanol largely elicits unique responses at distinct developmental stages spanning early embryogenesis.

**Fig 3.**
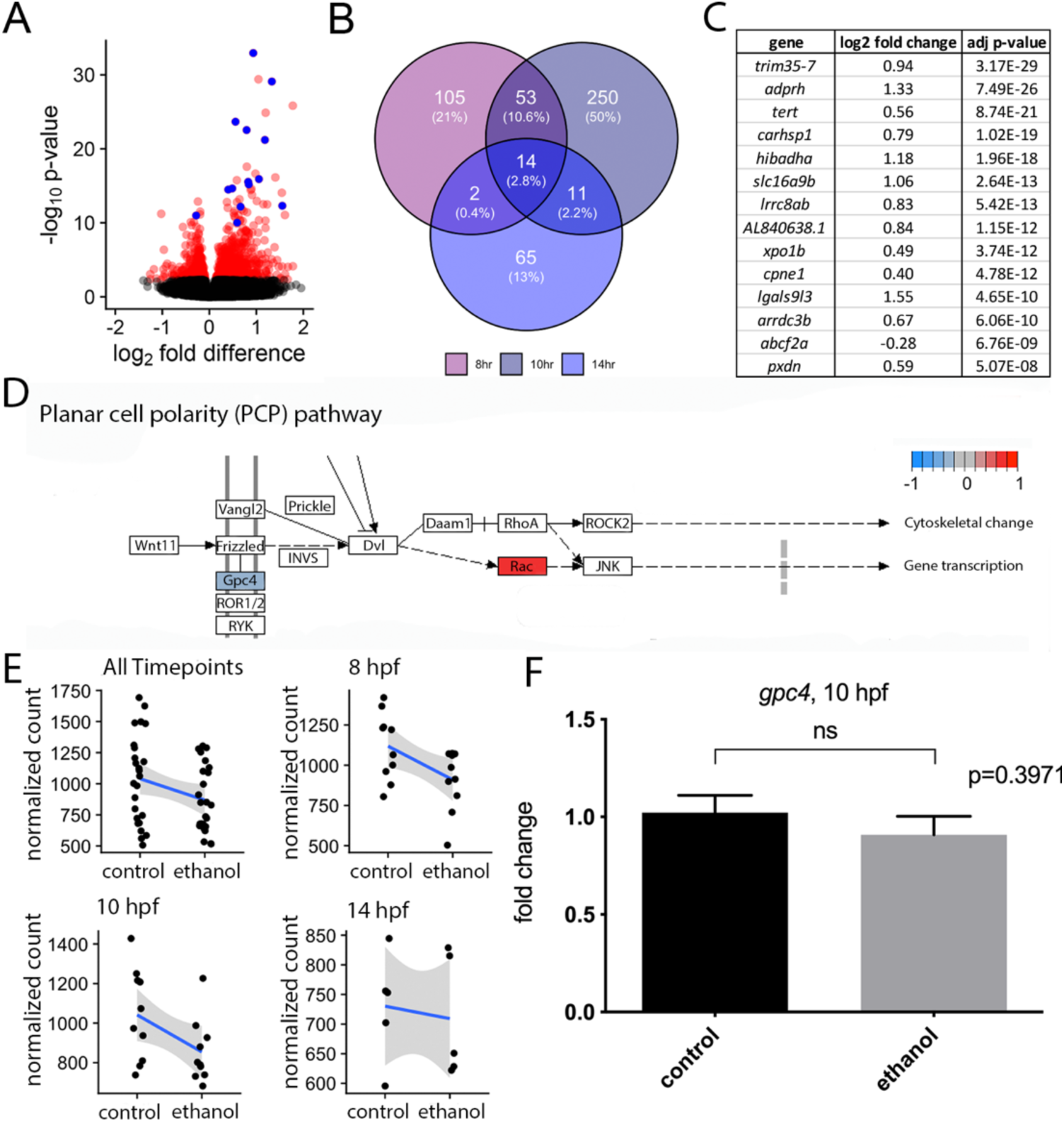
Effects on transcription are largely distinct between developmental time points. (A) Volcano plot showing differential expression due to ethanol treatment across all samples. Significant genes (FDR < 0.1)* are indicated in red. Genes that were significant at all time points (overlap) are indicated in blue. (B) Venn-diagram showing overlap of significant genes between the three individual timepoints. (C) Table of genes that were significant for each timepoint individually. (D) KEGG pathway schematic illustrating differential expression due to ethanol treatment in the Wnt/planar cell polarity (Wnt/PCP) pathway. Color coding indicates log_2_ fold differences due to ethanol treatment across all samples, red indicates upregulation due to ethanol, blue indicates downregulation due to ethanol. Only pathway members with significant changes were color coded. In the event that multiple genes from the dataset were annotated with the same pathway member, the log_2_ fold difference for the gene with the greatest absolute value for the difference was used.

### Ethanol does not affect the Wnt/PCP pathway at the transcriptional level

While *vangl2* mutants do not typically exhibit cyclopia, cyclopia is observed in double mutants between *vangl2* and other Wnt/PCP pathway members [15, 19]. One potential mechanism for the interaction between ethanol and *vangl2* would be the transcriptional misregulation of other Wnt/PCP pathway members. However, transcription of Wnt/PCP pathway members was largely unaffected by ethanol exposure. No Wnt/PCP pathway members were among the 14 shared DEGs across all timepoints (Fig 3C). KEGG pathway enrichment analysis further confirmed that ethanol had little effect on the Wnt/PCP pathway at the level of transcription (Fig 3D). Only two Wnt/PCP pathway members were moderately affected: ethanol exposure moderately decreased expression of the cofactor, *glypican 4* (*gpc4*), (log_2_ fold= −0.237; p-value = 0.036) across all timepoints [15, 20] and upregulated expression of *rac3a* (log_2_ fold = 0.737; p-value = 8.89E-06), a member of the Rho family of small GTPases [21]. We plotted the normalized read counts across each time point from the RNA-seq experiment to further investigate the dynamics of the potential alteration in *gpc4* levels. The expression of *gpc4* was modestly reduced in ethanol-treated embryos across the RNA-seq dataset (Fig 3E). Ethanol exposure consistently downregulated *gpc4* at 8 and 10 hpf, but the magnitude of the downregulation was relatively modest. To statistically compare ethanol-treated and control embryos, expression of *gpc4* at 10 hpf was investigated using qRT-PCR. This result demonstrated that *gpc4* was not significantly affected by ethanol exposure (Fig 3F). Thus, direct transcriptional alteration to the Wnt/PCP pathway is unlikely to explain the ethanol-induced phenotypes in *vangl2* mutants and heterozygotes.

### Modules of co-regulated genes related to ethanol exposure

To determine if ethanol disrupted networks of genes that could explain the *vangl2*- ethanol interaction, we next performed Weighted Gene Co-expression Network Analysis (WGCNA) [22]. This unsupervised network analysis identifies groups of genes, termed modules, based on correlated expression patterns across the samples. Modules are summarized by the first principal component for the expression estimates of the included genes, termed the module eigengene, which can be correlated with sample traits to identify biological significance. The cluster dendrogram generated in this analysis illustrates the presence of highly distinct and clustered modules (S3A Fig). Merging of similar modules produced eleven total modules (S3B Fig). Consistent with our PCA analysis, the module eigengenes were primarily correlated with time. We used these modules to further validate our ethanol exposure regimen.

Previous work has shown that ethanol delays development in a dose-dependent manner at concentrations equal to or greater than 1.5% ethanol [23]. For the experiments herein, ethanol-treated samples were morphologically stage-matched to control samples to exclude differences due to developmental age or delay. To confirm that the ethanol samples were indeed age-matched to the control samples at the level of transcription, we compared expression patterns of developmentally regulated genes between the ethanol and control samples from each timepoint. For developmentally regulated genes, we used the gene with the highest module membership (i.e. the hub gene) from each of the WGCNA modules that was associated with time (p < 0.05) (S4 Fig). Consistent with the results from the PCA (Fig 2C), the slope of the expression levels for each gene were similar in control and ethanol-treated samples across age, most clearly demonstrated in the magenta4, thistle1, and blueviolet modules (S4B,C,G Fig). Interestingly, we find a time-specific difference in the expression of hub genes in the honeydew1, greenyellow, and saddlebrown modules at 10 hpf (S4F,H,K Fig). Together, these data indicate samples were accurately age-matched and the observed changes were biologically relevant.

Two modules (mediumpurple4 and darkolivegreen4) correlated with ethanol treatment (S3B-D Fig). The purple and green modules positively and negatively correlated with ethanol treatment, respectively (S3C Fig). A significant differentially expressed gene from the green (S3G Fig) and purple module (S3H Fig) were selected for independent validation on independent biological replicate samples derived from the same wild-type zebrafish line using quantitative real-time RT-PCR (qRT-PCR). These results indicate that our RNA-seq faithfully represents transcript levels. The green module is only weakly downregulated and is enriched for genes encoding zinc finger (ZnF) proteins, which included *znf1015* (S3E Fig). This module is almost entirely composed of genes on chromosome 4, that have been shown to be co-regulated during these stages of development [24, 25]. The purple module revealed GO enrichment of transmembrane transporters, which included *slc16a9a* (S3F Fig). As an upregulated module enriched in transporters, these genes are likely to represent the physiological response to ethanol. Thus, while we did identify two modules associated with ethanol (S1 Table), the module membership did not provide clues to the interaction between ethanol and *vangl2*.

### Cyclopamine is predicted to mimic the effects of ethanol

One challenge in RNA-seq analyses is inferring the mechanism underlying a diseased or environmentally-perturbed state from a large set of differentially expressed genes. Individual functional analyses of significant gene-ethanol interactions are time consuming and thus inefficient. To circumvent this problem, we adopted a bioinformatics approach, utilizing the Library of Integrated Network-Based Cellular Signatures (LINCS L1000) toolkit [26]. This transcriptomic dataset includes gene expression data from nine human cancer cell lines exposed to thousands of small molecule drugs [26]. We queried the top 100 and 150 up- and down-regulated genes induced by ethanol exposure against the LINCS L1000 dataset using the clue.io platform (https://clue.io). The query generated a list of small molecules predicted to have a positive or negative correlation to the input signature (*i.e.* small molecules predicted to either mimic or antagonize the transcriptional effects of ethanol). We were particularly interested in those chemicals with positive correlation to ethanol as they would give insight into the mechanism of ethanol teratogenicity and highlight potential co-factors that exacerbate ethanol teratogenicity.

Interestingly, cyclopamine, a hallmark Shh pathway inhibitor that inhibits the core Shh pathway protein Smoothened (Smo), was predicted to positively correlate with the ethanol signature. The Shh pathway is critical for midfacial development and mice deficient in *Sonic Hedgehog* (*Shh*) exhibit severe brain and face malformations, including holoprosencephaly and a single medial eye (cyclopia) [27]. In zebrafish, reduction of *shh* or null mutations in *smo* similarly results in severe loss of craniofacial midline structures (i.e. the anterior neurocranium) [28]. Animal models have demonstrated ethanol is an environmental risk factor for holoprosencephaly, resulting in a characteristic set of midfacial defects, a hypomorphic forebrain, and in severe cases, cyclopia [16, 29–32]. Thus, attenuation of the Shh pathway could mechanistically explain the *vangl2*-ethanol interaction.

### Ethanol indirectly attenuates Shh signaling

To investigate the interaction of cyclopamine and ethanol, we first exposed wild-type zebrafish embryos to cyclopamine (50 μM) at shield stage (6 hpf) for 24 hours, mimicking the ethanol exposure window for the *vangl2* mutants. Embryos were fixed at 4 dpf and the cartilage and bone were stained with Alcian blue and Alizarin red, respectively. We observed a range of midfacial defects with the most severe phenotype being a complete loss of the anterior neurocranium and reduced spacing between the eyes (Fig 4A). Since cyclopamine was predicted to mimic the effects of ethanol, we next combined this low dose of cyclopamine with the subteratogenic dose of ethanol. Strikingly, all embryos presented with synophthalmia or cyclopia and significant reductions and defects of the cartilages of the neuro- and viscerocranium. To quantify this combinatorial effect, we measured the distance between the lenses and observed a significant reduction in co-exposed embryos relative to those exposed to either ethanol or cyclopamine alone (p<0.0001), suggesting a strong synergistic interaction (Fig 4B).

**Fig 4.**
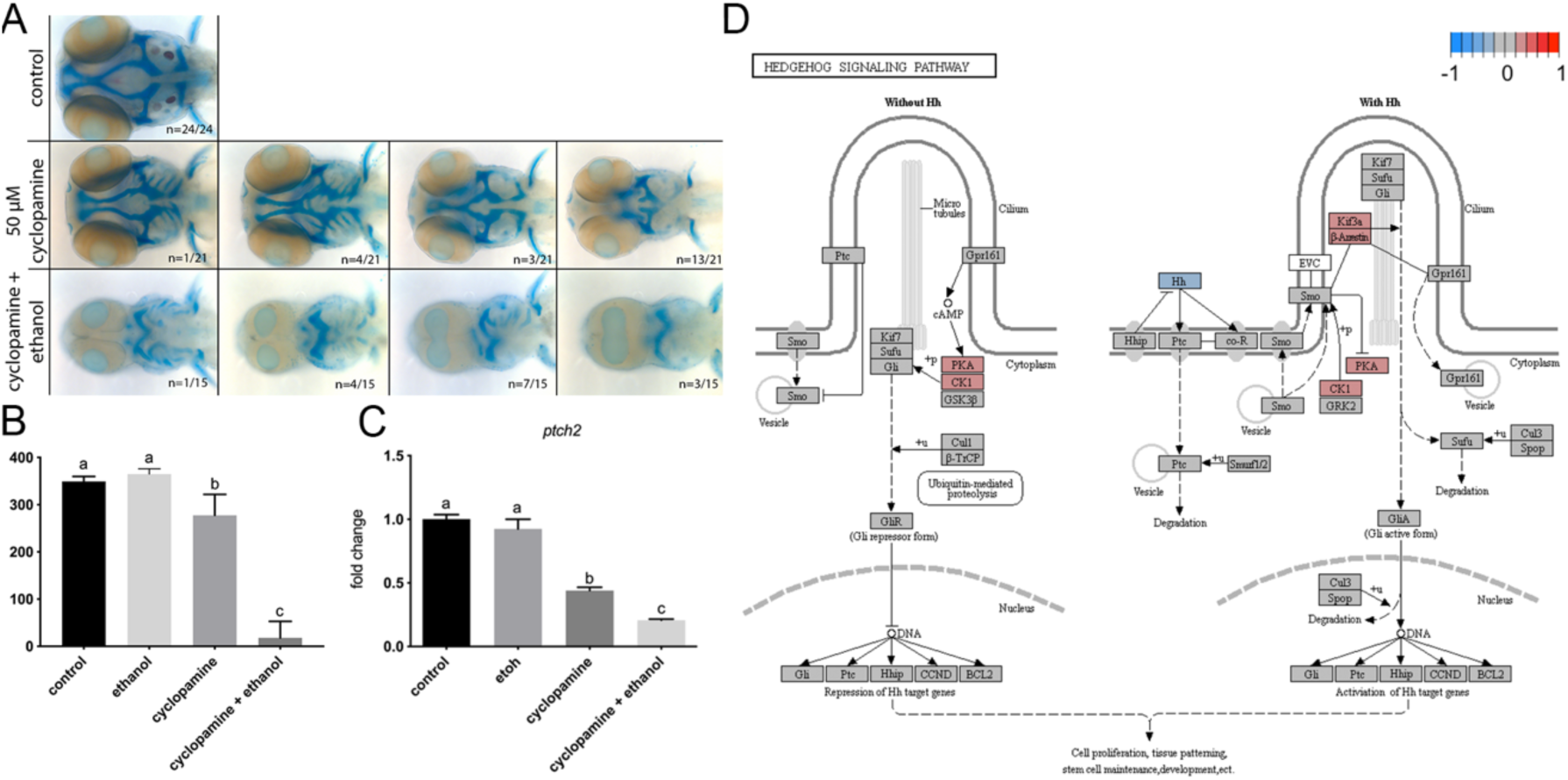
Ethanol indirectly attenuates Shh signaling. (A) Alcian blue and Alizarin red whole mount staining of untreated (control), cyclopamine-treated (50 μM), and ethanol- and cyclopamine-treated (1% ethanol plus 50 μM cyclopamine) wild-type embryos from 6 hpf to 4 dpf. Panels represent the spectrum of phenotypes observed for the treatment groups. Embryos were fixed at 4 dpf. Dorsal view, anterior to the left. (B) Quantification of the effect of ethanol and cyclopamine on the eye field. Inner lens-to-lens width was used as a morphometric measure of cyclopia. Both cyclopamine alone and cyclopamine and ethanol, were significantly different from controls (p<0.0001). (C) qRT-PCR of *ptch2* in 10 hpf wild-type embryos. *slc25a5* (*solute carrier family 25*) was used as a normalization control. (D) KEGG pathway schematic illustrating differential expression due to ethanol treatment in the sonic hedgehog (Shh) signaling pathway. Color coding indicates log_2_ fold differences due to ethanol treatment across all samples; red indicates upregulation due to ethanol, blue indicates downregulation due to ethanol. In the event that multiple genes from the dataset were annotated with the same pathway member, the log_2_ fold difference for the gene with the greatest absolute value for the difference was used.

In chick, ethanol exposure during somitogenesis has been proposed to suppress Shh signaling and induce apoptosis in cranial neural crest cells that make up the craniofacial skeleton [33]. However, work in zebrafish shows only a modest increase in cell death within the eye field at a much higher dose of 2% ethanol [34]. To ensure that cells in the eye field are not simply undergoing apoptosis, we performed a TUNEL cell death assay in ethanol-treated *vangl2* mutants at 11 hpf, prior to optic vesicle evagination. As expected, we failed to detect an increase in apoptotic cells within the eye field in *vangl2* mutants or their siblings (S5 Fig). Thus, Shh-mediated pro-survival signals do not appear to be significantly disrupted by ethanol.

To test for a direct effect of ethanol on Shh signaling, we quantified the relative expression of *ptch2*, a canonical read-out of Shh pathway activity, at bud stage (10 hpf). Subteratogenic doses of ethanol had no effect on the expression levels of *ptch2*, consistent with RNA-seq results (Fig 4D). While cyclopamine significantly reduced Shh signaling (p = 0.0006), ethanol did not further reduce *ptch2* levels significantly (p = 0.1115). KEGG analysis from the RNA-seq confirmed that ethanol does not affect the Shh pathway as a whole, nor does it reduce the levels of Shh target genes (i.e. *gli*, *ptch*) (Fig 4E). Thus, at normally subteratogenic doses, ethanol does not appear to directly attenuate Shh signaling.

### Ethanol disrupts convergent extension

The ethanol-induced *vangl2* mutant phenotype closely mirrors those in compound mutants between *vangl2* and other Wnt/PCP pathway members [15, 19]. These double mutants display a further reduction in convergent extension, as evidenced by a shorter and broader body axis [15]. Based on these data, we hypothesized that ethanol disrupts convergent extension, which would mislocalize the Shh signal and result in eye defects.

We performed *in situ* hybridization on untreated and ethanol-treated *vangl2* embryos to examine convergent extension and to test the hypothesis that the Shh expression domain is mislocalized in ethanol-exposed *vangl2* mutants. We analyzed expression of *shh* in the axial mesoderm and *paired box 2a* (*pax2a*) in the midbrain-hindbrain boundary at bud stage (10 hpf) [35]. Ethanol was initiated at high stage (3.3 hpf), when *vangl2* heterozygotes and homozygotes are equally sensitive to the effects of ethanol. We observed a gene and ethanol-dose dependent reduction in the length of the *shh* expression domain (extension) and an increase in the width of the *pax2a* expression domain (convergence) (Fig 5A).

**Fig 5.**
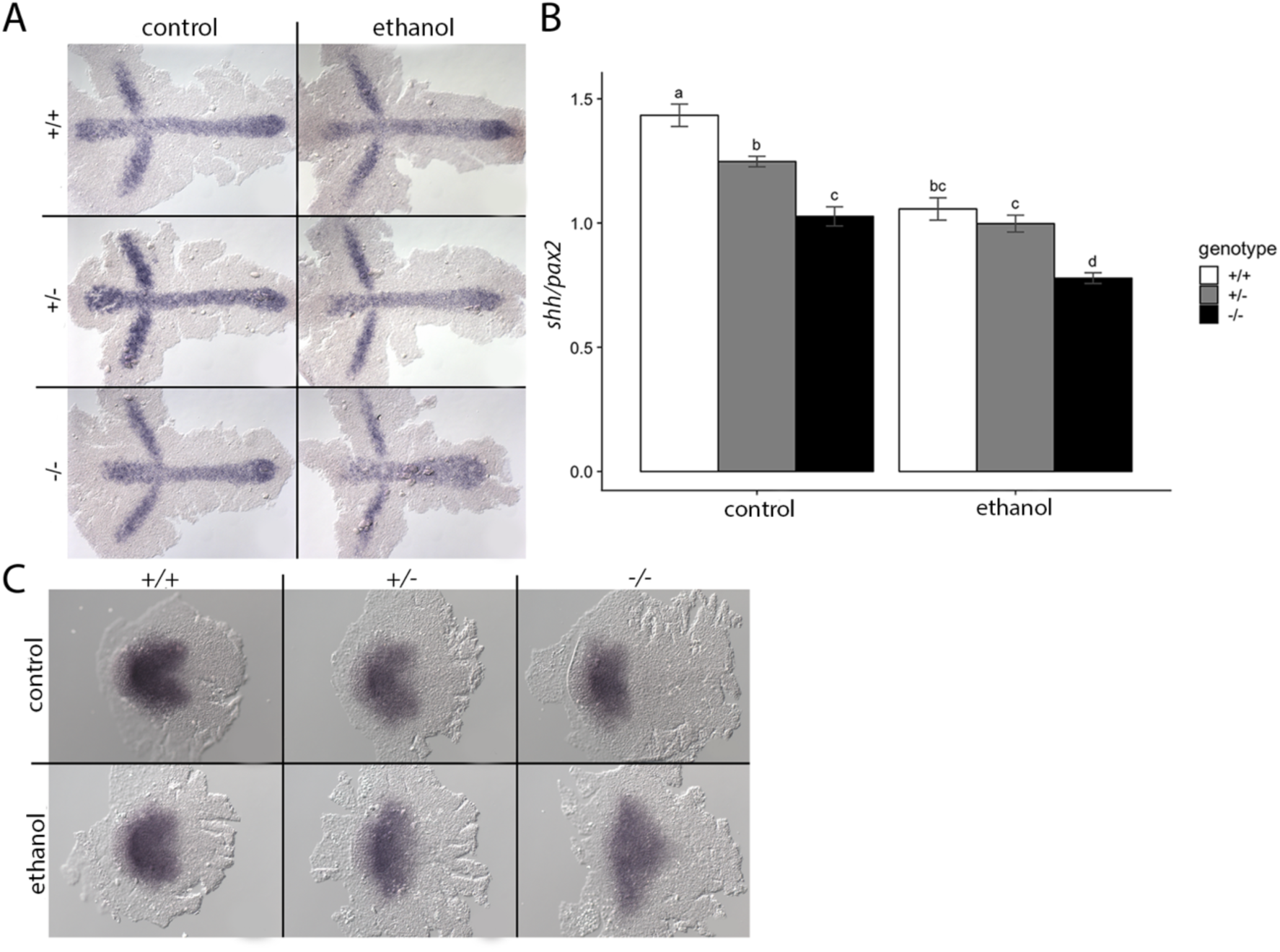
Ethanol disrupts convergent extension. (A) *vangl2* embryos were treated with 1% ethanol from 3.3 to 10 hpf. Embryos were subsequently harvested at 10 hpf for *in situ* hybridization with *shha* (midline) and *pax2a* (midbrain-hindbrain boundary) probes. Dorsal view, anterior to the left. (B) Quantification for the normalized *shh/pax2a* expression domains were calculated using the AxiovisionLE software. Two-way ANOVA followed by Tukey’s post-hoc test was used to analyze the results. Differences were observed between control genotypes and ethanol-treated homozygous mutants. (C) Expression pattern of transcription factor *six3a*, stained using whole mount *in situ* hybridization at 11 hpf.

To quantify the effect of ethanol on convergent extension, we plotted the normalized expression values of *shh*/*pax2* (Fig 5B). Post-hoc analyses (Tukey’s) revealed significant differences between untreated embryos across *vangl2* genotypes, indicating that *vangl2* gene dosage affects convergent extension. Similarly, we found significant differences between ethanol-treated mutants and their heterozygous and wild-type siblings (p = 0.017). This observation confirms that loss of *vangl2* results in reduced convergent extension movements, as evidenced by their short body axis. This phenotype was exacerbated with ethanol treatment, resulting in reduced convergent extension for *vangl2* heterozygotes and homozygotes, compared to their untreated counterparts. While there was a trend, we did not observe a difference in convergent extension between ethanol-treated *vangl2* heterozygotes and their wild-type siblings in their *shh/pax2* ratio. However, synophthalmia occurred in ethanol-treated heterozygotes but not wild-type embryos, demonstrating that the nonsignificant reduction in convergent extension in the heterozygotes can have a phenotypic consequence.

### Ethanol alters *six3* and *rx3* expression in the eye field

Transcription factors involved in the specification of the eye field have also been implicated in the mechanism of eye field separation. The expression of three of these transcription factors, *six3*, *rx3*, and *rx1*, are altered by ethanol exposure [34]. We examined the effect of ethanol on *six3a* and *rx3* in ethanol-treated *vangl2* mutants at the initiation of optic vesicle evagination. *In situ* hybridization of *six3a* in 11 hpf embryos shows a heart-shaped expression pattern in the prospective forebrain. The caudal indentation marks the splitting of the eye field into bilateral domains (Fig 5C). This expression pattern becomes more diffuse with loss of *vangl2*. In untreated homozygous mutants, we observed a shortening along the anterior-posterior (AP) axis and a broadening along the mediolateral axis at 11 hpf, consistent with the convergent extension defect. This expression pattern was further exacerbated in ethanol-treated mutants with complete loss of the caudal indentation. We observed a similar effect of genotype and ethanol on *rx3* expression at mid-evagination (12 hpf) (S6 Fig). At this stage, *rx3* is localized to the prospective forebrain and retina [36]. Ethanol-treated *vangl2* homozygous mutants exhibit a compressed expression domain, clearly displaying a reduction in convergent extension. Our expression analyses suggest the eye field is specified but mis-localized and fails to separate into bilateral domains, likely due to defects in mesodermal migration.

### Mutation in *gpc4* enhances cyclopia in a dose-dependent manner

Our data support a model in which ethanol interacts with *vangl2* via a combinatorial disruption of convergent extension. If ethanol disrupts convergent extension, which leads to interactions with *vangl2*, then further genetic disruption to convergent extension should exacerbate the ethanol-*vangl2* phenotype. Embryos deficient in *gpc4* similarly have a defect in convergent extension as evidenced by their shortened, broadened body axis [15]. Previous work in zebrafish has shown a functional interaction between *vangl2* and *gpc4*, where *vangl2;gpc4* double mutants were invariably cyclopic [15]. We conducted additional functional analyses to further examine the relationship between these two genes in the context of ethanol exposure. Consistent with Marlow *et al.*, double mutants were fully penetrant for cyclopia with or without ethanol. While 1% ethanol exposure altered the facial morphology of *gpc4* mutants (Fig 6A), it did not cause cyclopia. However, ethanol concentrations greater than 1%, which did not cause cyclopia in wild-type siblings, resulted in cyclopia in *gpc4* mutants (S7 Fig) [15]. Additionally, embryos carrying 3 mutant alleles (either *vangl2^-/-^;gpc4^+/-^* or *vangl2^+/-^;gpc4^-/-^*) were greatly sensitized to cyclopia when exposed to 1% ethanol (Fig 6B). Collectively these data indicate that in sensitized genotypes, ethanol disrupts convergent extension, thus mispositioning a source of Shh that is essential for separation of the eye field into bilateral primordia.

**Fig 6.**
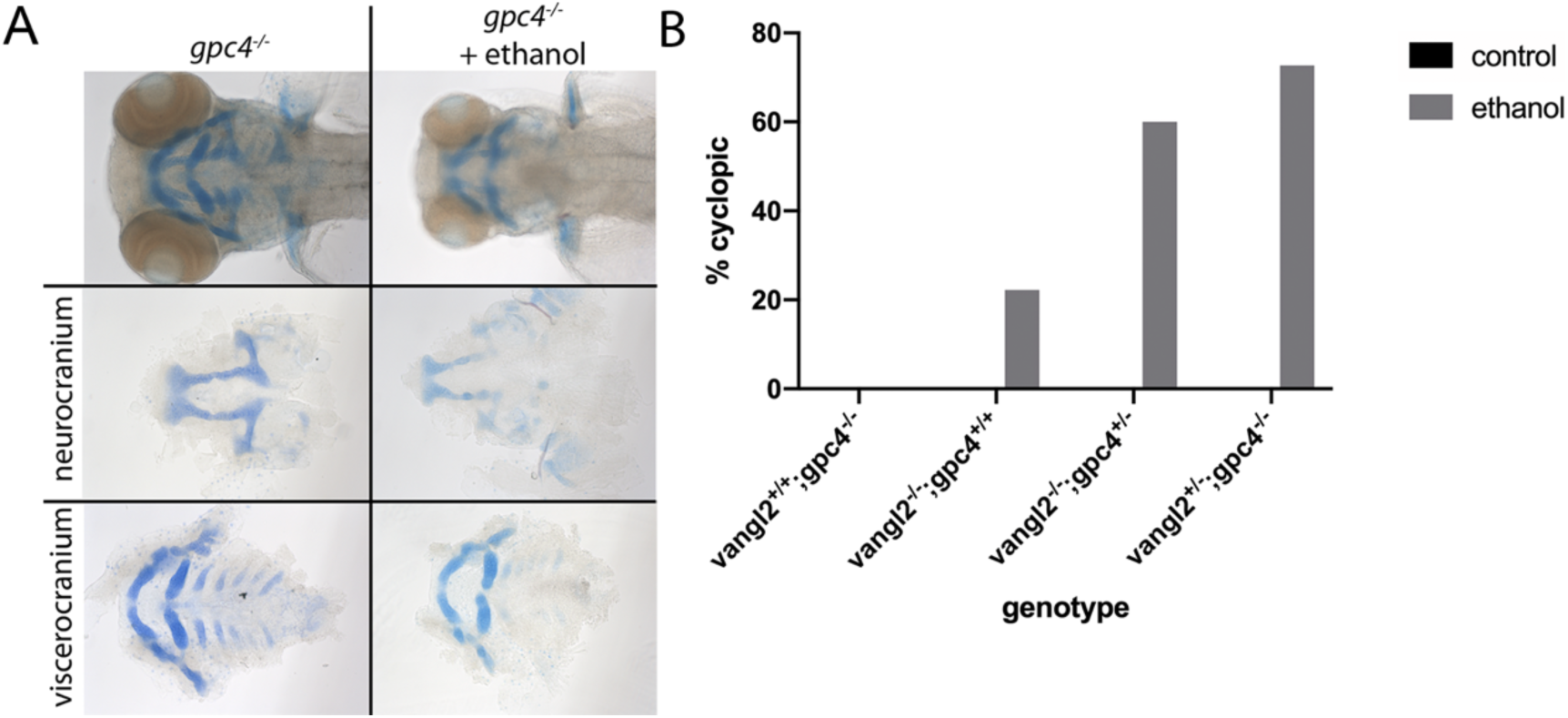
Mutation in *gpc4* exacerbates cyclopia in a dose-dependent manner. (A) Normalized read counts indicating expression of *gpc4* across different subsets of the dataset. (B) qRT-PCR of *gpc4* in 10 hpf wild-type embryos. *actb1* (*actin, beta 1*), *lsm12b* (*like-Sm protein 12 homolog b*), and *slc25a5* (*solute carrier family 25*) were used for normalization. (C) Alcian blue and Alizarin red whole- and flat-mount staining of untreated and 1% ethanol-treated (6 hpf – 4 dpf) *gpc4* homozygous mutants. Embryos fixed at 4 dpf. Dorsal view, anterior to the left. (D) Functional analyses of *vangl2;gpc4* double mutants. Enhanced cyclopia was observed in ethanol-treated *vangl2;gpc4* double mutants with loss of at least one copy of either gene.

### Blebbistatin phenocopies the ethanol-induced defect in *vangl2* mutants

If ethanol interacts with the Wnt/PCP pathway via convergent extension, then replacing ethanol with an inhibitor that disrupts cell movements, should similarly result in synophthalmia and/or cyclopia. We extended our analyses to test the effect of blebbistatin, a myosin II inhibitor that reduces axis elongation, in *vangl2* mutants [37]. Blebbistatin treatment during gastrulation (6-10 hpf) induced synophthalmia in *vangl2* heterozygotes and homozygotes (Fig 7A). We then quantified this interaction by measuring the inner lens-to-lens width and found a significant reduction in blebbistatin-treated *vangl2* homozygotes (Fig 7B). Furthermore, 26.92% and 100% of blebbistatin-treated heterozygotes and homozygotes, respectively, were cyclopic (Fig 7C). These data suggest ethanol further inhibits convergent extension in the sensitized background, *vangl2*, which not only disrupts their ability to elongate their body axis, but also inhibits the splitting of the eye field.

**Fig 7.**
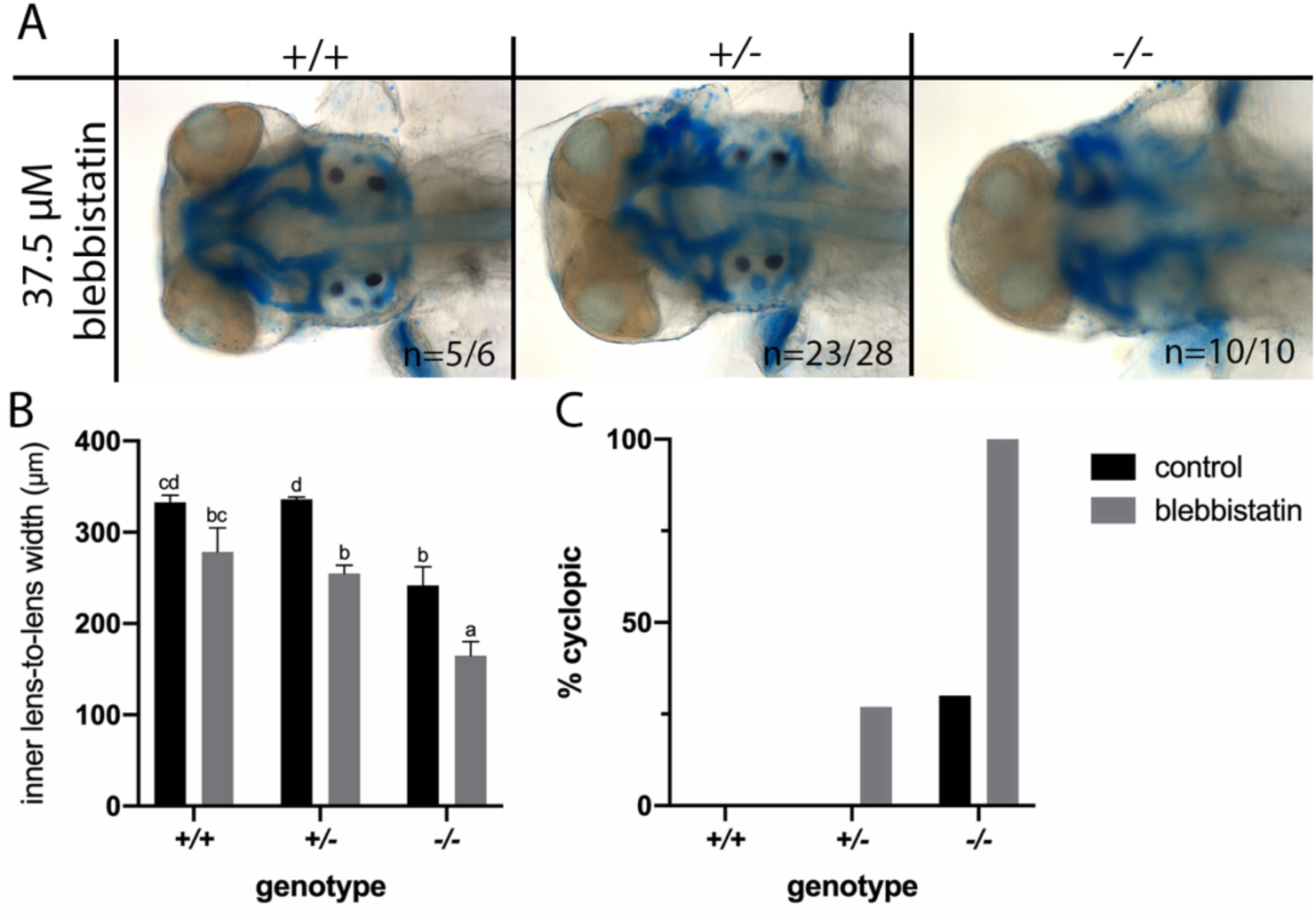
Blebbistatin phenocopies ethanol treatment in *vangl2* mutant embryos. (A) Alcian blue and Alizarin red whole-mount staining of *vangl2* embryos treated with 37.5μM blebbistatin (6-10 hpf). Embryos fixed at 4 dpf. Dorsal view, anterior to the left. (B) Quantification of the inner lens-to-lens width of control and blebbistatin-treated embryos. Lettering indicates differences in significance based on multiple comparisons test using Tukey’s HSD. (C) Representation of the percentage of control and blebbistatin-treated embryos exhibiting cyclopia. Genotypes indicated on the x-axes.

### Ethanol affects *vangl2* filopodia organization on migrating cells

Based on our data, cell dynamics underlying convergent extension are likely involved in the *vangl2*-ethanol interaction. During gastrulation, mesodermal cells adopt a bipolar shape and orient their actin-based cytoskeletal projections on their leading and trailing edge, with respect to the notochord, allowing for proper convergent extension [38]. In *vangl2* mutant embryos, these gastrula cells fail to elongate and align with the primary axis [39]. Furthermore, *vangl2* mutant ectodermal cells possess a greater number of filopodial protrusions than wild-type siblings and display projections in an unpolarized manner [40].

To determine whether ethanol may be exacerbating this phenotype, we analyzed filopodia protrusion length and orientation around the circumference of migrating live mesodermal cells. We injected memGFP mRNA into a single blastomere of *vangl2* embryos at the 8-32 cell stage, to obtain mosaic expression. We then treated a subset of the injected embryos with 1% ethanol from 6-10 hpf. Homozygous mutants were identified based on their reduced body axis elongation at 10 hpf. Cells were imaged at 40-60° from the primary axis. Spike-like filopodia were highly unpolarized in ethanol-treated *vangl2* mutants compared to their wild-type and heterozygous siblings (Figure 8A). We quantified the polarity and number of filopodia for each cell, with respect to the head (90°), the primary axis/notochord (180°), and the tail (270°) (Figure 8B). Rose diagrams suggest ethanol-treated *vangl2* mutants have more un-polarized projections on the anterior/posterior edge (red) as opposed to the leading (green) or trailing (blue) edge. We next reduced the number of bins in the rose diagrams to 90° and noticed a clear over- representation of filopodia in *vangl2* mutants on their anterior/posterior edge (Figure 8C). Tukey’s post-hoc analyses confirmed ethanol-treated *vangl2* mutants had significantly more filopodia on their anterior/posterior edge compared to other axes and their wild-type or heterozygous siblings (Figure 8D). These data confirm that ethanol exposure is associated with an increase in the number of filopodial projections oriented in the wrong direction, or perpendicular to the path of dorsal migration, in ethanol-treated *vangl2* mutants.

**Fig 8.**
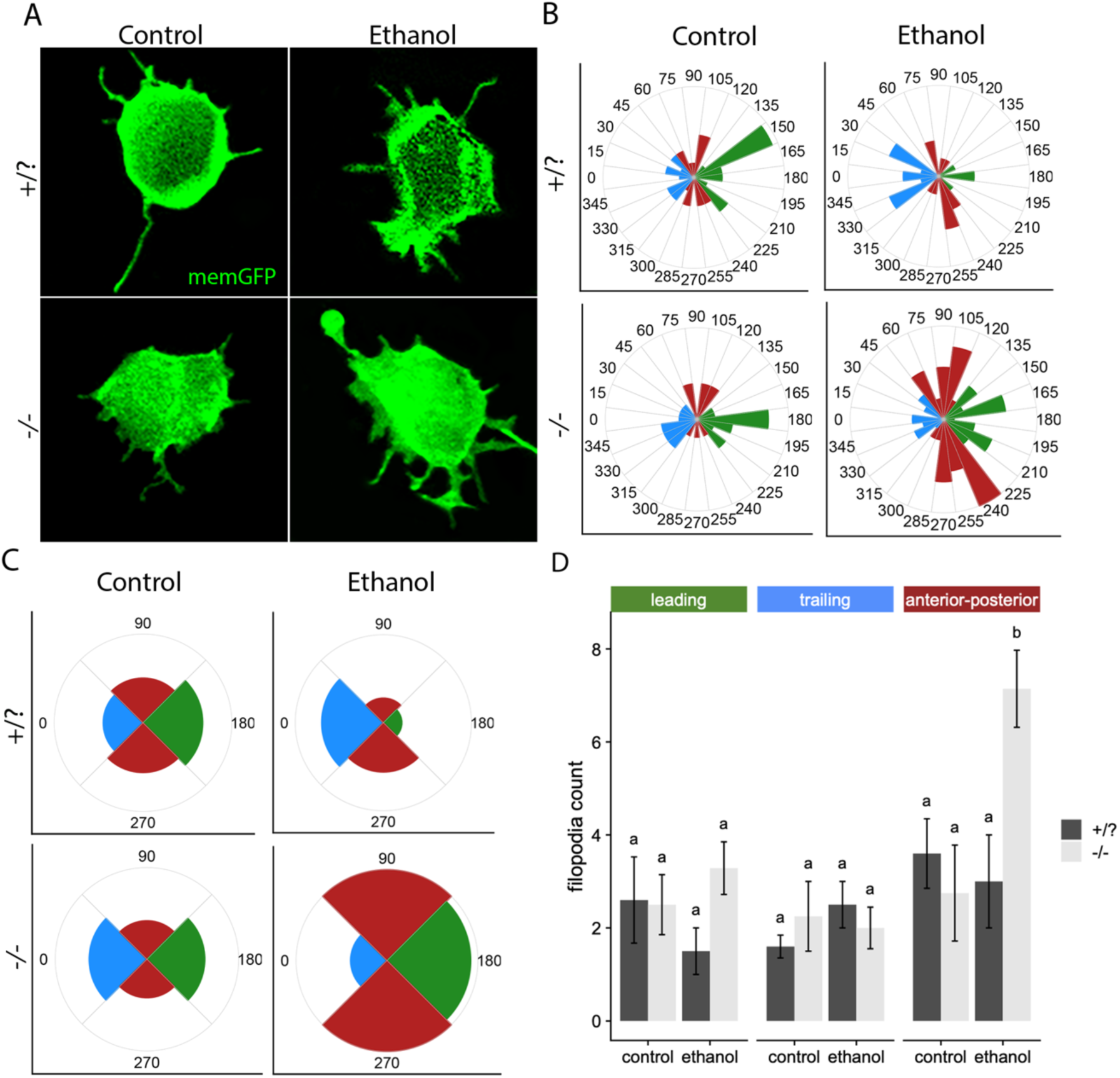
Membrane protrusions in untreated and ethanol-treated (6-10 hpf) *vangl2* embryos. (A) Confocal images of untreated and ethanol-treated *vangl2* mesodermal cells expressing memGFP at 10 hpf. n=6 (B) Rose diagram of filopodial projections representing length and distribution around the circumference of a cell. 24 bins, divided into 15o angles (C) Rose diagram of filopodial projections representing length and distribution around the circumference of a cell. 4 bins, divided into 90° angles. Horizontal plane (0°) represents the mediolateral axis; vertical plane (90°) represents the anterior-posterior axis.

## Discussion

### Molecular staging for the early zebrafish embryo

Despite their importance, the nature and mechanisms of gene-environment interactions are largely unknown. Here, we performed single embryo bioinformatic analyses to shed light upon the mechanism behind gene-environment interactions. A growing number of researchers are using the zebrafish for toxicological and/or gene-environment studies. One potential confounding variable in these types of studies is developmental delay. Our WGCNA analyses are rich in gene modules that similarly vary over developmental time, providing a way to molecularly stage embryos. Thus, our results will provide important resources to ensure the rigor and reproducibility of these analyses.

### The effects of ethanol on the transcriptome is subtle

Our hierarchical clustering analysis demonstrates that the effect of ethanol on the transcriptome is minor, within any given time point ethanol-exposed and unexposed embryos intermingled. PCA results support this, demonstrating that only 3% of the variance in the transcriptomic data is attributable to ethanol exposure. In sum, we found 1,414 differentially regulated genes across all time points with the majority of these genes being upregulated. Most differentially expressed genes were specific to a single time point. At any given time point there were between 92 and 328 differentially expressed genes. Thus, exposure to a normally subteratogenic dose of ethanol results in subtle and dynamic alterations to the transcriptome.

Our findings identified dramatically fewer differentially expressed genes than those observed in transcriptomic analyses of ethanol-exposed chicken and mouse embryos. In chicken 3,422 differentially regulated genes were observed after 6 hours of ethanol treatment and the preponderance (1,924 genes) were downregulated [41]. In a mouse microarray, 2,906 differentially expressed genes were identified 3 hours after ethanol injection [42]. Species-specific developmental trajectories or the tissue analyzed, cranial neural fold (chicken), headfold (mouse), versus whole embryo (our study) may account for some of these differences. However, it is important to note that the chicken and mouse strains analyzed are highly sensitive to the dose of ethanol used while our zebrafish AB strain is not (without the addition of a sensitizing mutation). Thus, the chicken and mouse data likely represent transcriptomic changes representative of pathogenesis. Our zebrafish data reflect a direct effect of ethanol, which may or may not be pathogenic depending upon the genetic and environmental milieu.

WGCNA analysis identified two modules (mediumpurple4 and darkolivegreen4) of genes that are coordinately altered by ethanol exposure. The vast majority of genes that negatively correlate with ethanol exposure (membership in darkolivegreen4) encode zinc finger proteins (ZnF) located on chromosome 4. Ethanol has similarly been shown to downregulate *Zinc finger protein, subfamily 1A*, 4 in mouse fetuses exposed to ethanol during early development [43]. However, the functional significance of these changes in ZnF proteins remains unknown and they often have diverse binding affinities and functions. The long arm of zebrafish chromosome 4 (*Chr4q*) is typically heterochromatic (condensed) and lacking in protein-coding genes [44]. However, it is not late replicating until the end of gastrulation or bud stage (10 hpf) [25]. Previous work in zebrafish has found ZnF proteins on chromosome 4 to undergo robust expression from the initiation of zygotic transcription until mid-gastrula stage [24]. Since many chromosome 4 genes are downregulated across ethanol-treated individuals during gastrulation, ethanol may interfere with replication timing, blocking the early-to-late replication switch, or chromatin remodeling. Based on these findings, the differentially expressed genes may follow disrupted expression of ZnF proteins due to chromatin remodeling.

### Ethanol interacts with *vangl2* through the indirect effects on the Shh pathway

Because Vangl2 is a core member of the Wnt/PCP pathway, a simple explanation for ethanol-sensitivity in *vangl2* mutants would be that ethanol transcriptionally dysregulates the Wnt/PCP pathway. However, our RNA-seq results indicated that this is not the case. The expression of *gpc4* was the only member of the Wnt/PCP pathway that was downregulated in our RNA-seq dataset. Cyclopia is observed in *vangl2;gpc4* double mutants [15] and we show that heterozygosity for *gpc4* further sensitized *vangl2* mutants to ethanol. However, the downregulation of *gpc4* was quite modest and unlikely to have any substantial impact on the phenotype of ethanol-exposed *vangl2* mutant.

We used an unbiased approach, the LINCS L1000 dataset to identify pathways that might be disrupted by ethanol and generate cyclopia in *vangl2* mutants. One of the top matches to the effects of ethanol was cyclopamine, a Hh pathway inhibitor. Shh signaling is critically important for development of the midface across species [27, 45–47]. In zebrafish *shha* and *smo* mutants have cyclopia because Shh signaling to the diencephalon is critical for separation of the eye fields [48, 49]. Furthermore, cyclopia observed in Wnt/PCP pathway mutants is due to the mispositioning of this source of Shh [16, 19]. Thus, disruption of the Shh pathway could mechanistically explain the ethanol sensitivity of *vangl2* mutants.

The Shh pathway has been implicated in ethanol teratogenesis. However, there has been much controversy regarding whether this interaction is direct or not [8]. As predicted by LINCS, we find a synergistic interaction between ethanol and the Hh pathway inhibitor cyclopamine. However, both our RNA-seq and qPCR data demonstrate that, at this timing and dose, ethanol does not alter the expression of direct readouts of Shh activity. Such an alteration would be expected if ethanol interacted directly with core components of the pathway. As supported by our *in situ* analysis, we conclude that ethanol indirectly effects the Shh pathway by mislocalizing the source of the ligand.

### Ethanol disrupts convergent extension by disrupting polarized cellular protrusions

The phenotypes observed in ethanol-exposed *vangl2* mutants suggest that ethanol further disrupts convergent extension. Our findings are consistent with previous observations that high doses of ethanol can generate phenotypes consistent with a disruption of convergent extension in zebrafish and Xenopus [16, 50, 51]. More recently, zebrafish microarray analyses demonstrated that disrupted gastrulation movements due to ethanol exposure were caused by a reduction in *pchd18a* expression [52]. The expression of *pchd18a* is not altered in our dataset. However, Sarmah and colleagues used the ethanol-sensitive TL strain and exposed embryos to ethanol from 2 to 8 hpf. This later difference is particularly intriguing giving our finding that the time window of greatest sensitivity to ethanol was different for *vangl2* heterozygotes and mutants, with heterozygotes being most sensitive at 3.3 hpf (the earliest time we tested). This raises the intriguing possibility that the earlier time window of teratogenesis is due to a different mechanism.

We propose that ethanol leads to midfacial defects and cyclopia, not by dysregulating Wnt/PCP at the level of transcription, but indirectly through inhibition of convergent extension and mislocalization of important developmental regulators like *shha*. At the start of convergent extension, lateral gastrula cells migrate and converge to the dorsal region of the developing embryo, where they intercalate between neighboring cells to drive extension of the body axis [20, 53, 54]. As these cells begin their dorsal migration, they elongate along their mediolateral axis and polarize their actin-based cytoskeletal processes medially and laterally to drive intercalation [54, 55]. Convergent extension movements during gastrulation also alter the shape of the eye field, whereby diencephalic precursor cells articulate both medially and caudally [56]. We know from previous work that *vangl2* mutant lateral gastrula cells fail to elongate their mediolateral axis and have a slight anterior bias in their dorsal trajectory [39, 57]. In addition to their shape changes, *vangl2* mutant ectodermal cells have more filopodia than wildtype cells and defective polarization (i.e. less filopodia localized on the trailing edge), resulting in an indirect dorsal trajectory [40]. Using live confocal imaging coupled with memGFP expression, we examined the effect of ethanol on filopodia number and polarity in *vangl2* embryos. Interestingly, we did not find more filopodia in *vangl2* lateral mesodermal cells relative to wildtype cells. Despite this, we find that ethanol-treated *vangl2* mutant embryos have more filopodia on their anterior/posterior edge relative to their mediolateral edge. In support of our hypothesis, we demonstrate that ethanol disrupts protrusion polarity, convergent extension, and *shha* localization, with greater disruption observed in *vangl2* mutants.

## Materials and Methods

### Zebrafish (Danio rerio) Care and Use

Zebrafish were cared for using standard IACUC-approved protocols at the University of Texas at Austin. The wild-type AB strain was used for RNA-seq analysis. The *vangl2^m209^* allele, originally described as *tri^m209^* [39], was obtained from the Zebrafish International Resource Center (ZIRC) as reported [13]. The gpc4^*fr6*^ line was provided by Dr. Lila Solnica-Krezel. Adult fish were maintained on a 14h/10h light-dark cycle at 28.5°C. Embryos were collected and staged according to morphology and somite number [58].

### Chemical treatments

Embryos were treated with 1% ethanol diluted in embryo media. This subteratogenic dose mimics an acute (binge-like) alcohol exposure roughly equivalent to a blood alcohol concentration of 0.19% in humans. AB embryos were treated with 50 μM cyclopamine (Toronto Research Chemicals C988400), diluted in embryo media. The concentration of ethanol (vehicle) in the embryo media was controlled for between treatment groups and equals a final concentration of 0.5% in the EM. Blebbistatin (Sigma- Aldrich B0560) was dissolved in DMSO and diluted to a concentration of 37.5 μM in 3 mL of embryo media. Following chemical exposure, embryos were washed 3x in embryo media and then grown to 4 dpf before fixation.

### mRNA injection

memGFP cloned in the pCS2 vector was linearized by *NotI* restriction endonuclease. Synthetic mRNA was generated using the mMESSAGE mMACHINE™ SP6 Transcription Kit (Invitrogen, AM1340). Embryos were microinjected at 100pg between the 8-16 cell stage [40].

### Sample collection and RNA extraction

Single embryos were manually dechorionated and collected in a 1.75 mL microcentrifuge tube with 500 mL of TRIzol reagent (Life Technologies, 15596-026). Embryos were homogenized with a motorized pestle (VWR, 47747-370) and stored at −80°C until RNA extraction. Total RNA was processed according to the TRIzol RNA isolation protocol. Samples were re-suspended with 50 μL of nuclease-free water and subsequently purified using the RNA Clean & Concentrator kit (Zymo, R1018). The concentration of each sample was determined using a Nanodrop spectrophotometer. The quality of total RNA was analyzed with the Agilent BioAnalyzer to ensure that the RNA Integrity Number (RIN) was ≥ 8. Samples were submitted to the Genomic Sequencing and Analysis Facility (GSAF) at the University of Texas at Austin. The GSAF performed standard RNA-Seq library preparations with poly-A mRNA capture.

### RNA-seq data processing

Sequencing on the NextSeq 500 platform produced an average of 40.8 million ± 1.4 million (SE) raw paired end reads per sample. Adapter trimming was performed using Cutadapt with a minimum length of 25 bp [59]. Following adapter trimming, we retained an average of 40.0 ± 1.3 million (SE) reads per sample. Genome Reference Consortium Zebrafish Build 10 (GRCz10) for *D. rerio* was downloaded from Ensembl [60]. Trimmed reads were mapped to the reference using STAR [61]. Mean mapping efficiency was 78.2% ± 0.8% (SE). Following mapping PCR duplicates were removed using Picard (https://broadinstitute.github.io/picard/). Duplication rate was estimated at 85% ± 0.8% (SE). Sorting and conversion between SAM and BAM files was performed using samtools [62]. Reads mapping to annotated genes were counted with HTseq version 0.6.1p1 using the intersection nonempty mode [63]. The final number of reads mapped to annotated genes was on average 3.9 ± 0.2 million reads per sample. Detailed instructions and example commands for implementing the data processing steps described above are available on Github (https://github.com/grovesdixon/Drerio_early_ethanol_RNAseq).

### Differential expression analysis

Normalization and statistical analysis of read counts was performed using DESeq2 [64]. Factors included in the differential expression models were *ethanol treatment* (control, treated), *developmental timepoint* (8 hpf, 10 hpf, and 14 hpf), and *sequencing batch* (experiment 1 or experiment 2). Because none of the 6 hpf samples were treated with ethanol, these samples were not included. We tested for differential expression associated with ethanol treatment using likelihood ratio tests—comparing the model including all three factors to a reduced model that did not include ethanol treatment. To further examine stage-specific ethanol effects we split the samples by developmental timepoint and tested for ethanol effects within each group.

### Weighted Gene Correlation Network Analysis (WGCNA)

Gene expression data were further analyzed with Weighted Gene Correlation Network Analysis (WGCNA) [22]. For input into the analysis, we used variance stabilized counts generated using the rlog function in DESeq2 [64]. Genes that were not sequenced across sufficient samples (4320 in total) were removed using the goodSamplesGenes function in the WGCNA package. Because there were no ethanol treated samples for the six hour timepoint, the five samples from this timepoint were removed before further analysis. To ensure sufficient expression for correlation detection, genes were further filtered based on a base mean expression cutoff of 5. We controlled for batch effects using the ComBat function from the R package sva [65]. We selected a soft threshold of 15, where the scale free topology model fit surpassed 0.8. WGCNA was run with a minimum module size of 10. Following network analysis, we tested for GO enrichment within modules using Fisher’s exact tests.

### GO enrichment analysis

Enrichment of Gene Ontology (GO) terms for ethanol responsiveness was tested using two-tailed Mann-Whitney U-tests [66] followed by Benjamini-Hochberg procedure for false discovery correction [67]. The results were plotted as a dendrogram tracing hierarchical relationships between significant GO terms. The direction of enrichment (for upregulation or downregulation) was indicated by text color and significance of enrichment by font type. An advantage of this approach is that it does not require an arbitrary cutoff to provide counts of “significant” and “non-significant” genes as in typical enrichment tests.

### Quantitative Real-Time RT-PCR (qRT-PCR)

To validate our RNA-seq data, we selected two genes to test using qRT-PCR. Total RNA was reverse transcribed using SuperScript™ First-Strand Synthesis System for RT-PCR (Invitrogen) with oligo-d(T) primers. qRT-PCR was performed with Power Sybr Green PCR Master Mix (Thermo Fisher Scientific, 4367659) on the Applied Biosystems ViiA™ 7 Real-Time PCR System. QuantStudio Real-Time PCR Software was used for data analysis using the 2^−ΔΔCt^ method. The endogenous control *lsm12b* was selected based on its stable expression profiles across treatment and stage groups in the RNA-seq datasets.

### KEGG enrichment methods and results

We tested for enrichment of significant upregulated and downregulated genes among KEGG pathways separately using Fisher’s exact tests. For each KEGG pathway, we tested a two-way contingency table with inclusion in the KEGG pathway as columns, and significant (FDR < 0.05) upregulation or downregulation as rows. P-values were then adjusted for multiple tests using Benjamini Hochberg Method [67].

Three KEGG pathways were enriched for significant differential expression. Two for upregulated genes, and one for downregulated genes. The upregulated KEGG pathways (dre00250 and dre00480) were (1) Alanine, aspartate and glutamate metabolism, and (2) Glutathione metabolism. The downregulated KEGG pathway (dre03013) was RNA transport. The pathview figures for these are in kegg_pathways/significant_pathway_figures/ in the git repository.

### Cartilage and Bone Staining and Measurements

Embryos were fixed at 4 days post fertilization (dpf) and stained with Alcian blue for cartilage and Alizarin red for mineralized bone [68]. Whole-mounts of *vangl2^m209^* embryos were captured using a Zeiss Axio Imager A1 microscope. To assess the degree of cyclopia, the distance between the medial edges of the lenses was measured using the AxiovisionLE software.

### *In Situ* Hybridization

Antisense digoxygenin-labeled riboprobes for *shha*, *pax2*, *dlx3*, *six3a* and *rx3* (together as a gift from Dr. Steve Wilson) were used. Whole-mount *in situ* hybridization was performed as described [69]. Images were captured using the Zeiss Axio Imager A1 and expression domains were measured using the AxiovisionLE software. An ANOVA and post-hoc Tukey’s test were used for statistical analyses.

### TUNEL Staining

Whole-mount TUNEL staining was based on previous characterizations [12]. Samples were fixed overnight in 4% paraformaldehyde in PBS (PFA) at 4°C. Samples were dehydrated in methanol and subsequently rehydrated in phosphate-buffered saline containing 0.5% Triton X-100 (PBTx). Samples were permeabilized with 25 ug/mL proteinase K (1mg/ml) in PBT for 30 min. After two, 5 min washes with PBTx, samples were fixed with 4% PFA for 20 min at room temperature. Residual PFA was removed with four, 5 min washes of PBTx. Samples were incubated with 50 μl of 1:10 Enzyme:TUNEL reagent (TdT and fluorescein-dUTP) (Roche, Cat No. 11684795910) at 37°C for 3 h in the dark. The reaction was stopped with two, 5 min washes of PBTx. Confocal images were captured with a Zeiss LSM 710.

### Confocal Imaging

Control and ethanol-treated (6-10 hpf) embryos were dechorionated and staged at 10 hpf. Homozygous mutants were distinguished between wild-type and heterozygotes by their body axis elongation at bud stage. Live embryos were mounted in methyl cellulose. Images were acquired using a Zeiss LSM710 confocal microscope using a 10x and 60x objective.

### Quantification of filopodia

Number of filopodia was acquired through projections of Z-stacks using the 60x objective on the Zen software. Angle of projections were calculated using Fiji.

Rose diagrams of the mean number of filopodia found were plotted using geom_bar() and the coord_polar() functions from ggplot2. Mean numbers of filopodia for different embryonic regions were compared between groups using ANOVA and Tukey’s ‘Honest Significant Difference’ method. Significantly distinct means were assigned based on the Tukey’s ‘Honest Significant Difference’ results using the R package multcomp [70].

## ACKNOWLEDGEMENTS

We thank Dr. Steve Wilson for his kind contributions of the *six3a* and *rx3* riboprobes and Dr. John Wallingford for his contributions of the memGFP plasmid. We also thank Dr. Lila Solnica-Krezel for providing the *gpc4^fr6^* line. We are grateful to Anna Louise Percy, Angie Martinez, and Cadianna Cotham, for maintenance and care of all zebrafish lines. Lastly, we are grateful to our former undergraduate researchers, Jenna Beam, Elaine Avshman, and Jennyly Nguyen, for their assistance in the lab.

## Supporting Information

**S1 Fig.**
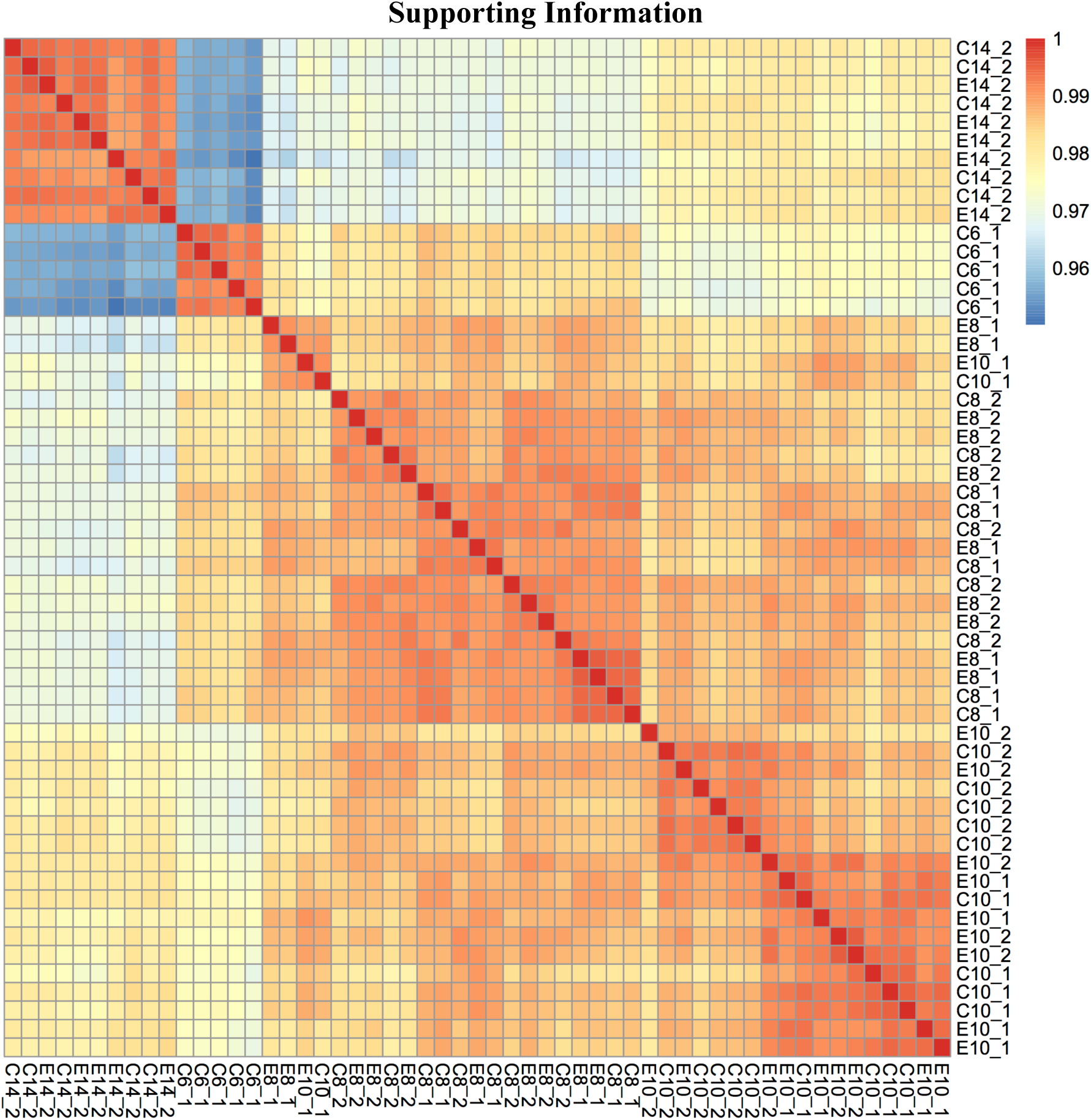
Time is the major driver of variation in the dataset. Heatmap showing overall correlation of gene expression among samples. Samples were hierarchically clustered based on similarity. Treatment [control or ethanol treated (C or E)] and developmental age [hours post fertilization (6, 10, 8, or 14)] are indicated in the row and column labels.

**S2 Fig.**
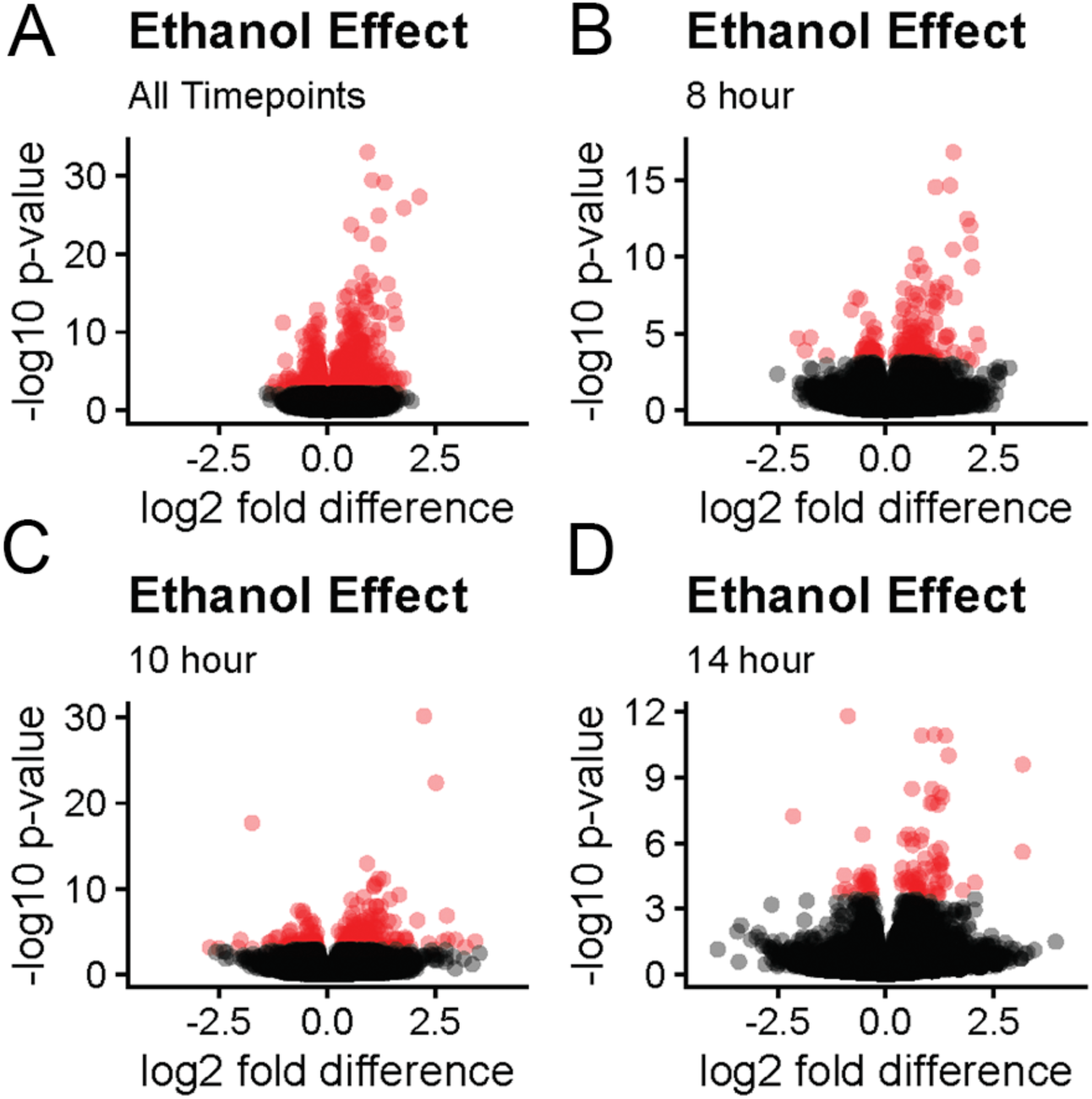
There are more upregulated than downregulated genes among ethanol-treated individuals. Volcano plot showing variation in the transcriptional response to ethanol treatment across developmental timepoints. Significant genes (FDR < 0.1) are indicated in red. For each subset, the names of the topmost significantly dysregulated genes are noted near gene’s data point (A) All timepoints combined together (B) 8 hpf only (C) 10 hpf only (D) 14 hpf only.

**S3 Fig.**
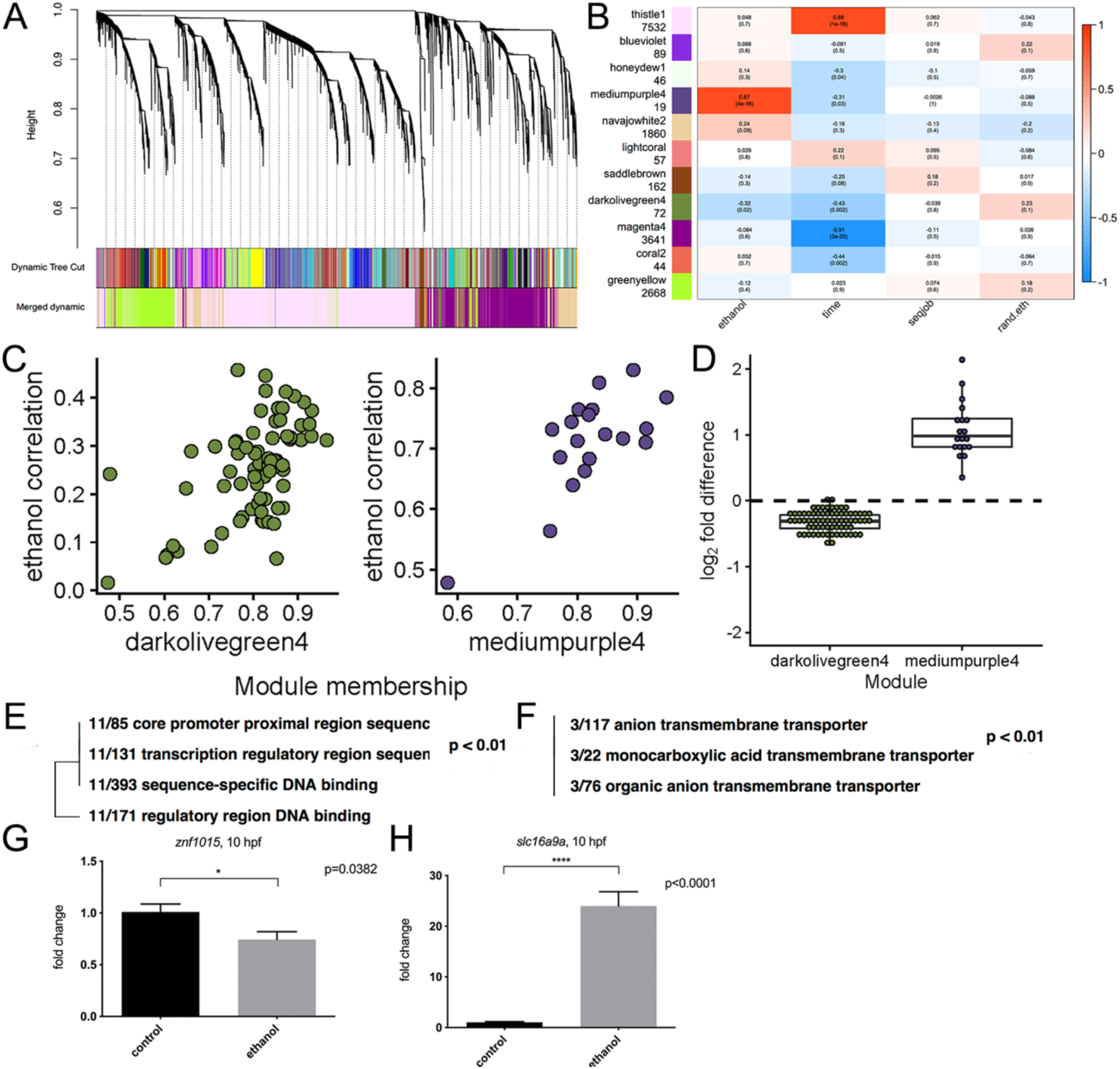
WGCNA identifies two modules that are significantly correlated with ethanol exposure. (A) Dendrogram illustrating the hierarchical clustering of the genes and with their corresponding modules colors. The top layer of colors indicates the Dynamic Tree Cutoff, including all assigned modules, before merging by module similarity. The bottom layer indicates the module colors after merging. These merged modules were used for further analysis. (B) Heatmap of module-trait correlations. The eigengene for each module was correlated with ethanol treatment (ethanol), hours post fertilization (age), experimental batch (seqjob), and as a negative control, a randomly shuffled version of the ethanol treatments (rand.eth). Intensity of the color in each cell indicates the strength of correlation between the module (row labels) and the sample trait (column labels). Two modules, (mediumpurple4 and darkolivegreen4) significantly correlated with ethanol treatment (p < 0.05). (C) Scatterplots of correlation with ethanol treatment against module membership. Each datapoint is a gene assigned to the indicated module. Ethanol correlation is the Pearson correlation between the gene’s expression level and ethanol treatment. Module membership is the correlation between the genes expression level and the module eigengene and describes how well the gene matches the overall patterns of the module. (D) Boxplot of log_2_ fold differences due to ethanol for the two significant modules. (E) Gene ontology enrichment tree for Molecular Function for the darkolivegreen4 module. (F) Gene ontology enrichment tree for Molecular Function for the mediumpurple4 module. (G) Changes in gene expression from the RNA-seq were validated using wild-type embryos at 10 hpf. *slc16a9a* was selected from the mediumpurple4 module and (H) *znf1015* was selected from the darkolivegreen4 module. Fold change indicates the degree of change between untreated control and ethanol-treated stage-matched embryos. *lsm12b* (*like-Sm protein 12 homolog b*) was used as a normalization control gene.

**S4 Fig.**
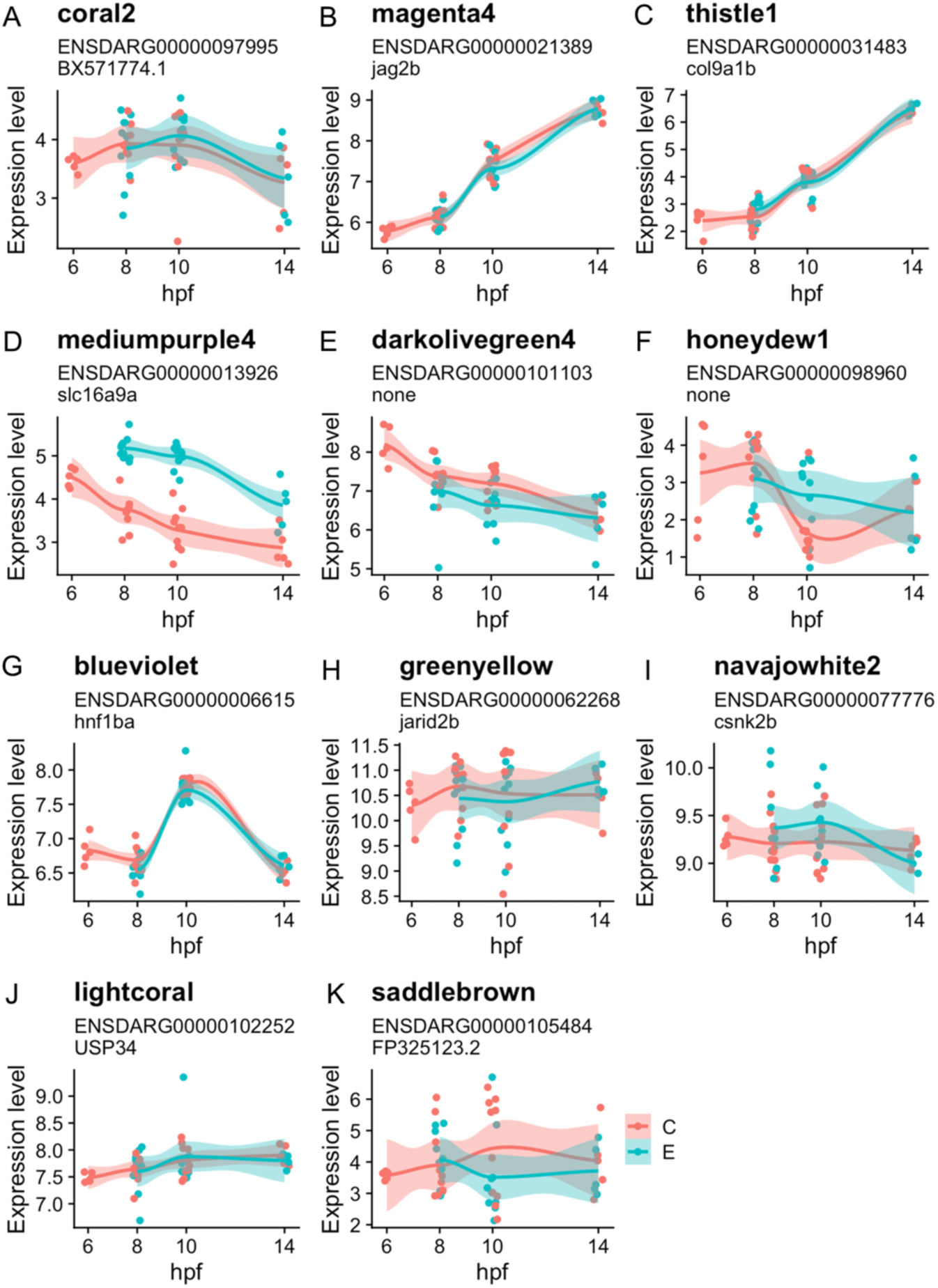
Expression of hub genes for development-related modules indicate ethanol did not retard developmental progression. (A-K) Development-related WGCNA modules were identified as those with significant relationship to hours post fertilization (Pearson correlation; p<0.05). The hub gene for each of these modules was identified as the gene assigned to that module with the highest module membership (defined as the correlation of the gene’s expression level with the module eigengene). Normalized expression levels for the hub genes were plotted against hours post fertilization. The relationship between expression and time is largely consistent between the ethanol treated (E, teal) and control (C, pink) samples. Even for the two modules associated with ethanol treatment (mediumpurple4 and darkolivegreen4), the slopes of the lines are very similar, indicating subteratogenic ethanol exposure did not cause significant developmental delay.

**S5 Fig.**
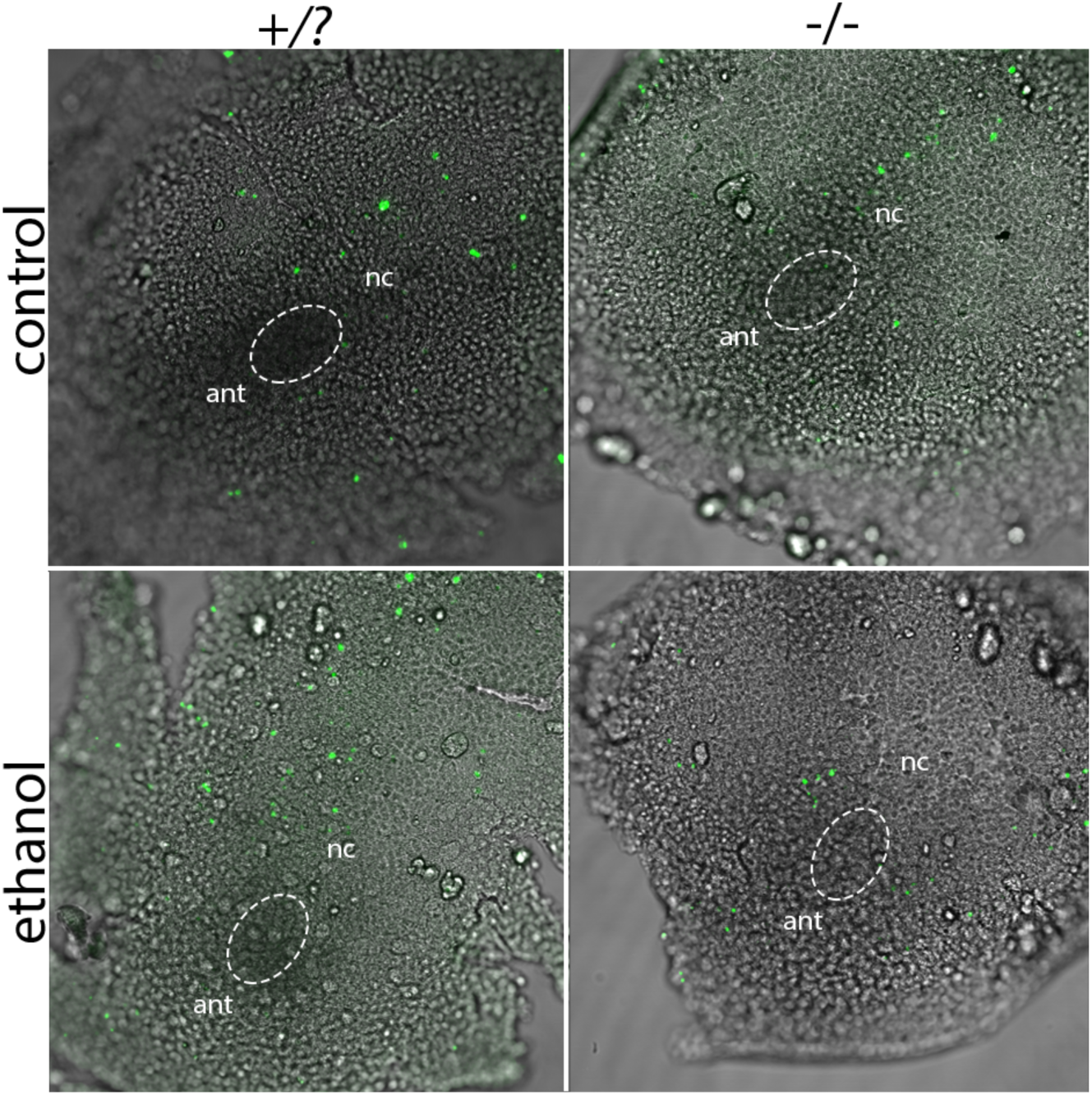
Cell death by TUNEL at 11 hpf in untreated and ethanol-treated *vangl2* mutants. The number of positive cells in the eye field (indicated by dashed circle) was not higher in homozygous or ethanol-treated mutants. ant = anterior; nc = notochord

**S6 Fig.**
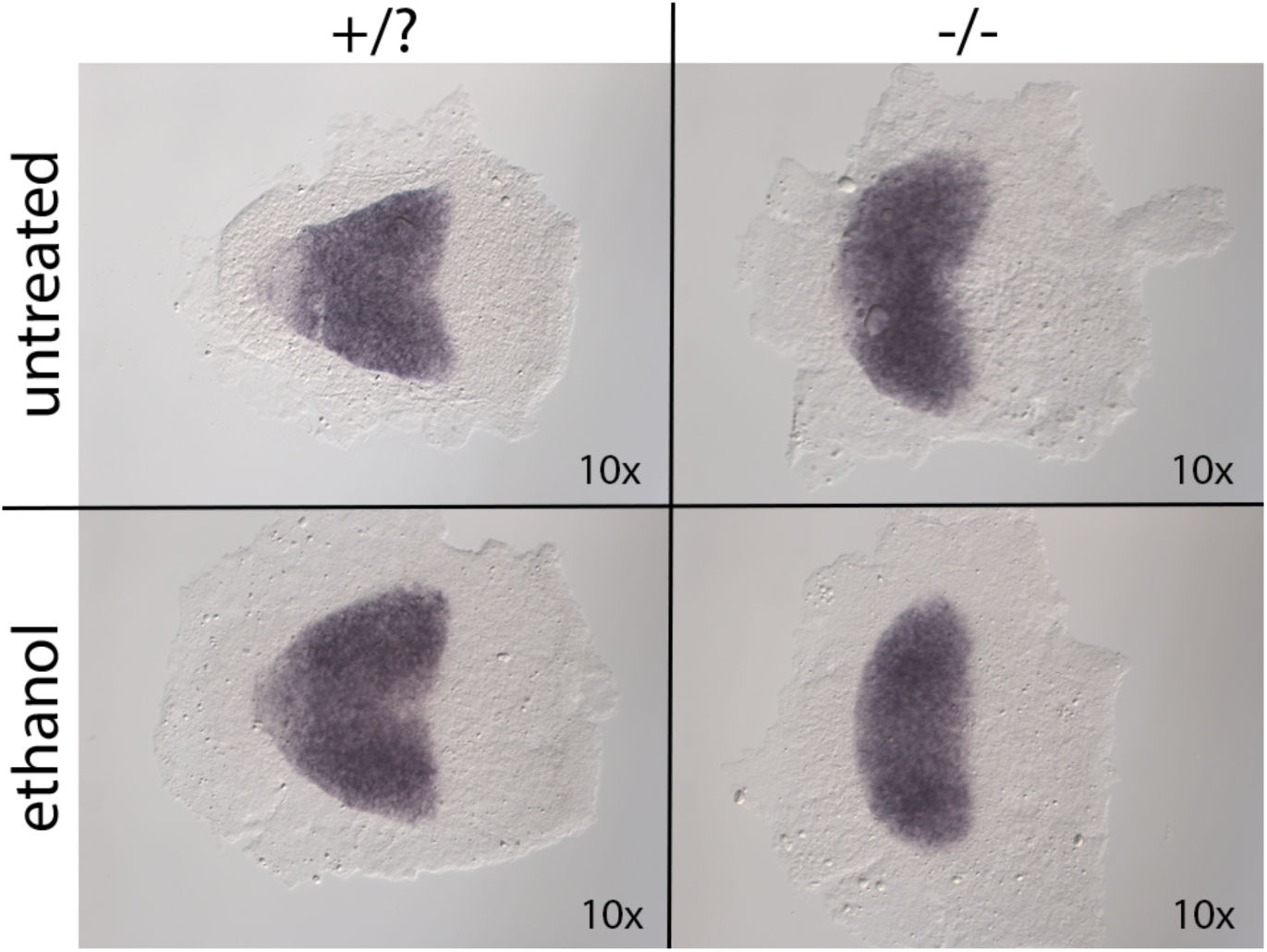
Ethanol alters *rx3* expression in the eye field. Expression pattern of transcription factor *rx3*, stained using whole mount *in situ* hybridization at 12 hpf. Dorsal view, anterior to the left.

**S7 Fig.**
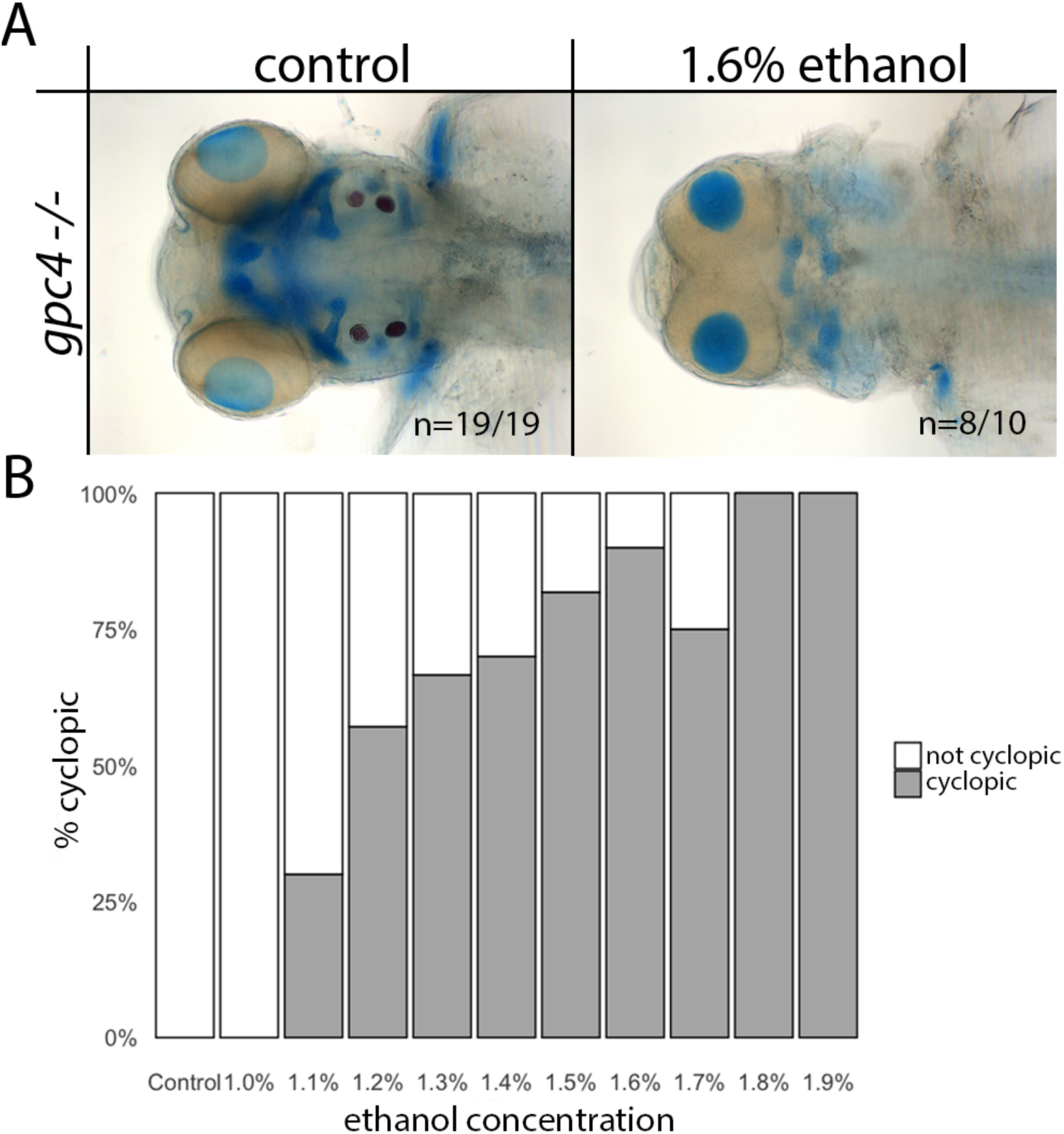
Ethanol interacts with *gpc4*. (A) Alcian blue and Alizarin red whole-mount staining of untreated and 1.6% ethanol-treated (6 hpf – 24 hpf) *gpc4* homozygous mutants. Embryos fixed at 4 dpf. Dorsal view, anterior to the left. (B) Dose-response curve of *gpc4* mutants treated with 1-1.9% ethanol (6-30 hpf).

